# SPTBN2 promotes an immunosuppressive tumor microenvironment and cross-resistance to anti-cancer therapies

**DOI:** 10.64898/2026.03.30.715365

**Authors:** Quoc Thang Bui, Raghavendra Basavaraja, Mayura R. Dhamdhere, Ågnes Holczbauer, Luca Paruzzo, Puneeth Guruprasad, Michael Scaglione, Yingying Tang, Yuchen Sun, Daniel P. Beiting, Erik K. Nash, Hossein Fazelinia, Lynn A. Spruce, Alice Wang, Kai Tan, Wei Guo, Crystal S. Conn, Yi Fan, Constantinos Koumenis, Vladimir S. Spiegelman, Hallgeir Rui, J. Alan Diehl, Matthew J. Atherton, Ben Z. Stanger, Will Bailis, Marco Ruella, Serge Y. Fuchs

**Affiliations:** Department of Biomedical Sciences, School of Veterinary Medicine, University of Pennsylvania, Philadelphia, PA 19104, USA; Dept. of Medicine, University of Pennsylvania, Philadelphia, PA 19104, USA; Department of Pathology and Laboratory Medicine, University of Pennsylvania, Philadelphia, PA 19104, USA; Department of Pathobiology, School of Veterinary Medicine, University of Pennsylvania, Philadelphia, PA 19104, USA; Proteomics Core Facility, Children’s Hospital of Philadelphia, Philadelphia, PA 19104, USA; Graduate Group in Genomics and Computational Biology, University of Pennsylvania, Philadelphia, PA, USA; Department of Pediatrics, University of Pennsylvania Perelman School of Medicine, Philadelphia, PA, USA; Dept. of Biology, University of Pennsylvania, Philadelphia, PA 19104, USA; Dept. of Radiation Oncology, University of Pennsylvania, Philadelphia, PA 19104, USA; Dept. of Pediatrics, Pennsylvania State University College of Medicine, Hershey, PA 17033, USA; Dept. of Pharmacology, Physiology and Cancer Biology, Thomas Jefferson University, Philadelphia, PA 19107, USA; Dept. of Biochemistry, Case Western Reserve University School of Medicine, Cleveland, OH 44106, USA; Department of Clinical Sciences and Advanced Medicine, School of Veterinary Medicine, University of Pennsylvania, Philadelphia, PA 19104, USA

**Keywords:** SPTBN2, immune suppression, immunotherapy, trogocytosis, memory T-cells, CAR T-cells, BTLA. tumor microenvironment, chemotherapy, cross-resistance

## Abstract

Immunosuppressive tumor microenvironment (TME) inactivates CD8+ cytotoxic lymphocytes (CTLs). Here, we identify SPTBN2 spectrin as a key immunosuppressive regulator induced in CTLs in response to nutritional deficit. In human pancreatic and colorectal cancers, SPTBN2 expression negatively correlated with CTL infiltration and patients’ survival. In TME of mouse pancreatic and colorectal adenocarcinomas, SPTBN2 inactivated intratumoral CTLs, stimulated tumor growth and conferred cross-resistance to anti-cancer therapies. SPTBN2 knockout protected CAR T-cells from trogocytosis and increased their memory state. SPTBN2 maintained levels of cell surface proteins such as BTLA that undermine CAR T-cell cytotoxicity and promote exhaustion. Re-expression of BTLA largely reversed phenotypes in SPTBN2-deficient CAR T-cells. In manufactured CAR T cells, SPTBN2 was associated with their clinical failure in pediatric patients with leukemia. Accordingly, ablation of SPTBN2 in CAR T-cells increased their cytotoxicity, in vivo persistence and therapeutic effects indicating that SPTBN2 can be targeted to increase the efficacy of anti-cancer therapies.

## INTRODUCTION

Immunosuppressive tumor microenvironment (TME) in solid tumors attenuates the anti-tumor immune responses, promotes tumor growth and progression, and undermines the efficacy of anti-cancer treatments ^1, 2^. At the foundation of these phenotypes is the inhibition of the anti-tumor CD8+ effector cytotoxic T lymphocytes (CTLs) by many factors such as deficit of oxygen and nutrients, immunosuppressive regulatory T cells, myeloid cells, and cancer-associated fibroblasts ^2-4^. The sheer redundancy of mediators undermining CTL inactivation complicates the efforts to target diverse TME elements. An alternative approach is to focus on identification, characterization, and targeting of key regulators that act within CTLs to suppress their anti-tumor effects ^2, 5^.

The surfaceome of CTLs contains numerous important ligands, receptors and regulators that can be engaged by the immunosuppressive mechanisms of the TME but also targeted by therapeutic agents ^6^. Global and specific regulators of localization and stability of cell surface proteins can define whether these CTLs will be capable of recognizing and eliminating malignant cells. Interfering with these mechanisms is successfully utilized by the solid tumors to generate immune privileged niches and establish immunosuppression that drives tumor growth, progression and resistance to therapy ^1, 2^.

Here, we report that the spectrin beta non-erythrocytic 2 (*SPTBN2*) gene acts as a key regulator of the immunosuppressive tumor microenvironment. *SPTBN2* appeared amongst the genes upregulated in response to nutritional stress in human and mouse CD8+ T-cells ^7^. *SPTBN2* encodes β-III-spectrin – a protein that plays a key role in linking integral membrane proteins with actin and the cellular cytoskeleton. Dominant mutations in this gene cause the neurodegenerative disease - spinocerebellar ataxia type 5 ^8^. Mice expressing analogous mutation or entirely lacking *Sptbn2* develop neurological defects, including ataxia and motor dyscoordination, that become evident around 6 months of age ^9, 10^.

Spectrins, including SPTBN2, regulate the localization and cell surface stability of numerous signaling proteins ^11, 12^. Given this important housekeeping role, it may seem surprising that tumors can co-opt the spectrin activities to further their growth and progression. However, current literature indeed suggests such roles for spectrins in cancer. For example, the β-II-spectrin (SPTBN1) contributes to the signaling and transcriptional responses driven by the transforming growth factor-β (TGF-β) and plays a key role in gastrointestinal and liver cancers ^13-15^. Likewise, in malignant cells, SPTBN2 acts on diverse cell surface proteins such as claudins and SLC7A11 cysteine channel and promote cell growth and survival in several cancer types ^16-19^. These studies provide rationale to develop therapeutics that could hamper abnormally increased SPTBN2 activities. However, the feasibility of utilizing such agents and their potential safety has not been established given the lack of information regarding the role of SPTBN2 in the non-malignant cells within the immunocompetent TME.

Here we report that SPTBN2 is induced in intratumoral CTLs and acts as a potent negative regulator of their anti-cancer activities. Expression of SPTBN2 in the TME reduced tumor infiltration by CTLs, undermined their expression of cytokines and effector molecules and upregulated markers of exhaustion. Accordingly, expression of SPTBN2 in the TME promoted tumor growth and resistance to chemotherapy and immune checkpoint inhibitors. Expression of SPTBN2 in the manufactured CAR T cells was associated with their clinical failure in pediatric patients with acute lymphocytic leukemia. Conversely, inactivation of SPTBN2 in the CAR T-cells promoted their memory differentiation, rendered them resistant to exhaustion, increased their persistence in vivo and augmented their therapeutic activities.

## RESULTS

### SPTBN2 is upregulated in the TME and drives intratumoral angiogenesis and stromagenesis

We have previously reported that depriving human and human CD8+ T-cells off key nutritional components elicits profound adaptive changes in transcriptional and translational gene expression profiles ^7^. Among affected genes, *SPTBN2* emerged as a candidate of interest because its total and ribosome-associated mRNA levels were significantly increased in murine (**Fig 1A**) and human (**Fig S1A**) CD8+ T-cells deprived of glucose or various amino acids. These results prompted us to investigate the putative roles of SPTBN2 in the metabolic adaptation of CD8+ CTLs to nutrient-deprived TME conditions and in functional modulation of these cells.

**Figure 1.**
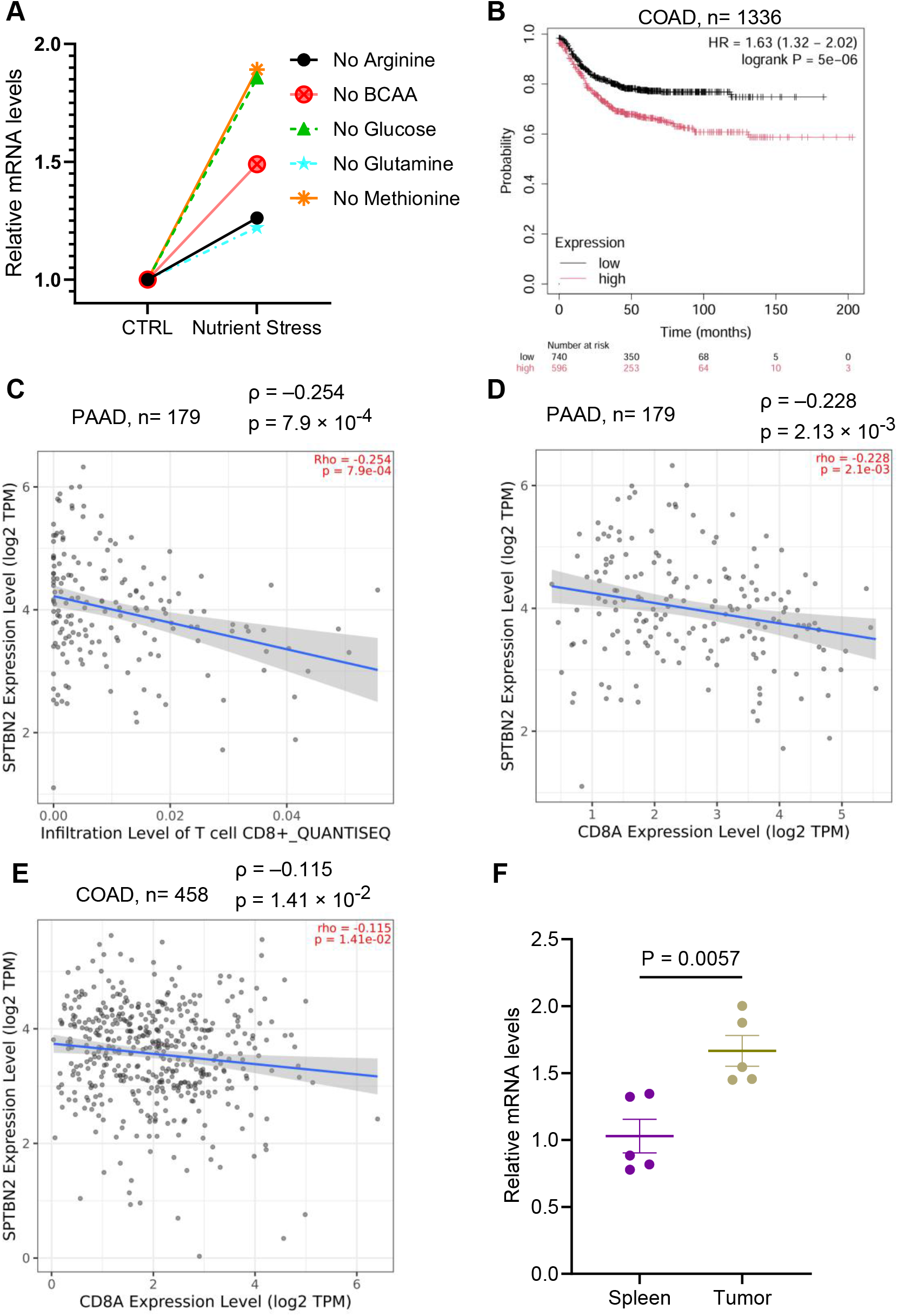
SPTBN2 is upregulated in the tumor microenvironment. A. Normalized levels of *SPTBN2* mRNA associated with polyribosomes in CD8^+^ T-cells cultured under conditions of deprivation of glucose, arginine, glutamine, methionine, or branched-chain amino acids (BCAA). Data is presented as a fold change relative to the control group (complete media); n=3. B. Association between *SPTBN2* expression and overall survival in human COAD (n=1336) samples, analyzed using the KMplot database. Patients were stratified into high and low expression groups based on the median SPTBN2 expression. The hazard ratio (HR), log-rank p-value, and 95% confidence interval (CI) are shown. C. Correlation of infiltration of CD8^+^ T cells and SPTBN2 expression in human PAAD (n=179) samples from TCGA, analyzed using TIMER 2.0. Statistical analysis was performed using the Spearman correlation test; the correlation coefficient (ρ) and p-value are indicated in the plot. D. Correlation of expression levels for *CD8A* and *SPTBN2* in human PAAD (179) samples from TCGA. Gene expression correlation was analyzed using TIMER2.0, and the Spearman correlation coefficient (ρ) and p-value are indicated. E. Correlation of expression levels for *CD8A* and *SPTBN2* inhuman COAD (n= 458) samples from TCGA, performed using TIMER2.0. The correlation coefficient (ρ) and corresponding p-value are shown. F. qPCR analysis of *Sptbn2* mRNA expression in T cells isolated from tumors or spleens from MC38 s.c. tumors bearing mice (n = 5). Expression levels were normalized to *Gapdh*. Data is presented as fold change relative to the control (spleen) group and shown as mean ± SEM. Statistical comparisons were conducted using the Student’s t-test.

Bioinformatic analysis of TCGA human cancer data linked the expression of *SPTBN2* with shorter overall survival for patients with colon (**Fig 1B**) and pancreatic (**Fig S1B** and ^19^) cancer. The pro-tumorigenic effects of SPTBN2 expression in malignant cells have been documented in several human malignancies ^17-21^. Here we focused on the importance of SPTBN2 in the cells of the TME.

We observed a negative association between SPTBN2 expression and infiltration of pancreatic adenocarcinoma tumors with CD8+ T-cells (**Fig 1C**). Further analyses demonstrated that expression of *SPTBN2* was inversely correlated with expression of *CD8A* gene, which is typically expressed by CTLs, in pancreatic (**Fig 1D**) and colorectal (**Fig 1E**) cancers. Similar negative associations between SPTBN2 and tumor-infiltrating CTLs were observed in lung adenocarcinoma (**Fig S1C**), lung squamous cell carcinoma (**Fig S1D**), and prostate adenocarcinoma (**Fig S1E**). Prompted by demonstration that SPTBN2 expression is induced in many types of tumors relative to normal tissue ^19^, we compared the levels of *Sptbn2* mRNA in mouse T-cells isolated from either MC38 colon adenocarcinoma tumors or from the spleens of tumor-bearing mice. We found that tumor-infiltrating T-cells express higher levels of *Sptbn2* compared to splenic T-cells (**Fig 1F**). Collectively, these data suggest that nutritional stress and perhaps other TME conditions upregulate SPTBN2 in tumor-infiltrating CTLs and indirectly implicate SPTBN2 in shaping the immunosuppressive TME.

To determine the importance of SPTBN2 in the non-malignant TME cells we used *Sptbn2*^-/-^ knockout mice (**Fig S2A**). These mice develop neurological defects including ataxia and motor discoordination starting at 6 months of age ^9^. We used these mice aged between 8-12 weeks when they did not display any overt phenotypes. At this age, the extensive immunophenotyping of splenic cells in naïve *Sptbn2*^-/-^ mice did not reveal any difference with WT littermates in numbers of CD4+ or CD8+ T-cells (**Fig S2B**), NK, dendritic cells, granulocytes or monocytes (**Fig S2C**). Furthermore, splenic *Sptbn2*-null CD8+ T-cells did not display statistically significant differences in the levels of activation (CD69), proliferation (Ki-67), exhaustion (PD-1, TIM-3 and LAG-3), or cytotoxic (IFN-γ, TNF-α or granzyme B) markers compared to WT counterparts (**Fig S2D**).

We inoculated WT or *Sptbn2*^-/-^ littermates with syngeneic MH6419c5 pancreatic ductal adenocarcinoma (PDAC) tumors and compared gene expression in the tumor tissues (**Fig 2A** and **Supplemental Table 1**). This comparison revealed a notable increase in diverse signatures/pathways including ion membrane transport, steroid and alcohol metabolism and others in tumors growing in the *Sptbn2*^-/-^ hosts (**Fig 2B**). These changes were consistent with a hypothesis that expression of SPTBN2 in the TME may affect both immune and non-immune stromal compartments of the tumors.

**Figure 2.**
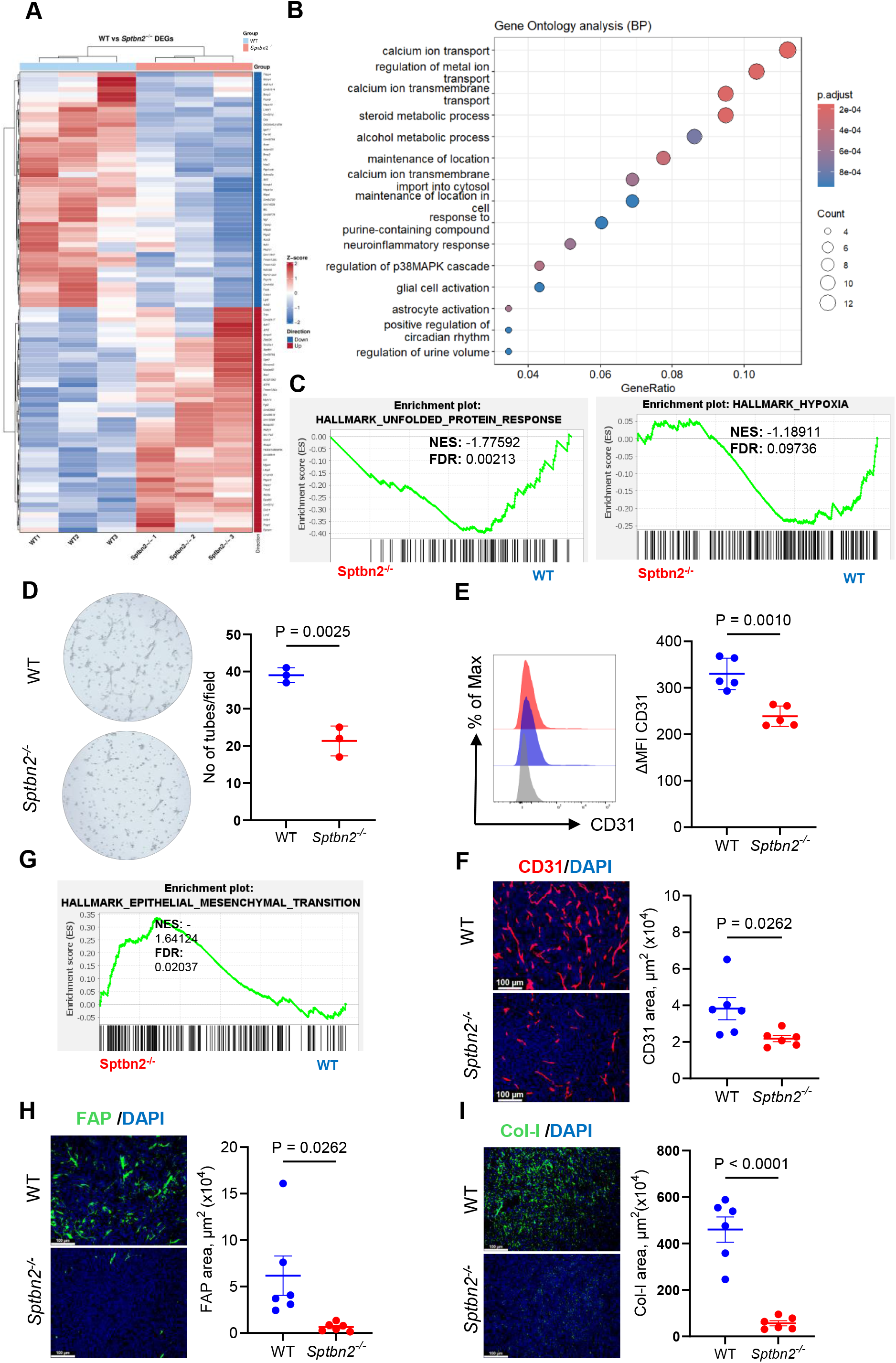
SPTBN2 supports intratumoral angiogenesis and stromagenesis. A. Heatmap of genes differentially expressed in s.c. “cold” MH6419c5 tumor tissues grown in WT and *Sptbn2*^*-/-*^ mice (n = 3). B. Gene ontology enrichment analysis of differentially expressed genes in MH6419c5 tumors growth on WT and *Sptbn2*^*-/-*^ mice (n = 3). C. Gene Set Enrichment Analysis (GSEA) of unfolded protein response and hypoxia hallmarks in MH6419c5 tumors grown in WT and *Sptbn2*^*-/-*^ mice. Normalized enrichment scores (NES) and false discovery rate (FDR) are indicated. D. Representative images (left) and quantification (right) of tube formation assays using lung-derived ECs from naïve WT and *Sptbn2*^*-/-*^ mice (n = 3). E. Flow cytometry analysis of CD31 on live cells isolated from subcutaneous MH6419c5 tumors growing in WT and *Sptbn2*^*-/-*^ mice (n = 5). F Representative immunofluorescence images of CD31 staining in orthotopic pancreatic MH6499c4 tumors growth in WT and *Sptbn2*^*-/-*^ mice (n=6), along with quantification of CD31-positive areas (scale bar: 100µm). G. GSEA of gene signature associated with epithelial-mesenchymal transition in MH6419c5 tumor growth in WT and *Sptbn2*^*-/-*^ mice. H. Representative immunofluorescence images and quantification of FAP staining in orthotopic pancreatic MH6499c4 tumor growth in WT and *Sptbn2*^*-/-*^ mice (n=6). Scale bar: 100µm. I. Representative immunofluorescence images and quantification of collagen-I staining in orthotopic pancreatic MH6499c4 tumor growth in WT and *Sptbn2*^*-/-*^ mice (n=6). Scale bar: 100µm. Data are presented as mean± SEM. Statistical comparisons were conducted using the Student’s t-test.

Intriguingly, we observed changes in gene signatures associated with hypoxia and unfolded protein response (**Fig 2C**) indirectly indicating potential involvement of SPTBN2 in tumor perfusion and prompting us to investigate the importance of SPTBN2 in angiogenesis. No visible vascular abnormalities were observed in naïve *Sptbn2*^-/-^ animals. Furthermore, normal pancreatic tissues did not show any difference in angiogenesis between WT and *Sptbn2*^-/-^ animals (**Fig S2E**). However, VEGF-activated lung endothelial cells (ECs) from *Sptbn2*^-/-^ mice exhibited deficient vascular tube formation compared with WT ECs *ex vivo* (**Fig 2D**). Accordingly, flow cytometry (**Fig 2E**) and immunohistochemistry (**Fig 2F**) analyses of orthotopic MH6449c4 PDAC tumor tissues revealed that *Sptbn2* is required for efficient tumor angiogenesis.

Furthermore, tumors grown in the *Sptbn2*^-/-^ animals exhibited an altered expression pattern of genes involved in the epithelial-mesenchymal transition **(Fig 2G**). This finding, together with reported role of a related SPTBN1 in the TGF-β pathway ^15^ prompted us to analyze intratumoral stromagenesis. Levels of fibroblast activation protein alpha (FAPα), a marker of cancer associated fibroblasts (CAFs), was found comparable in normal mouse pancreatic tissues from WT and SPTBN2 knockout mice (**Fig S2F**). However, MH6449c4 orthotopic PDAC tumors from the *Sptbn2*^-/-^ mice displayed significantly lesser levels of FAP (**Fig 2H**) as well as collagen I (**Fig 2I**) compared to tumors from WT mice. These results suggest that SPTBN2 in the TME contributes to intratumoral stromagenesis.

### SPTBN2 inactivates tumor-infiltrating CTLs, promotes tumor growth and confers cross-resistance to anti-cancer therapies

Given SPTBN2-associated expression of genes involved in the regulation of T-cell function, including IL-2 signaling, IFN-γ pathway, inflammatory responses and allograft rejection (**Fig 3A**), we used immunohistochemistry (**Fig 3B**) and flow cytometry (**Fig S3A**) analyses to characterize the intratumoral CD8+ T-cells from WT or *Sptbn2*^-/-^ mice. The MH6499c4 PDAC tumors from *Sptbn2*^-/-^ mice were characterized by an increased infiltration of CTLs (**Fig 3B**). Accordingly, knockout of SPTBN2 in the TME was associated with greater numbers of CTLs in MC38 tumors (**Fig 3C**). Importantly, SPTBN2-deficient CD8+ T-cells isolated from MC38 tumors displayed increased levels of cytotoxic cytokines and mediators such as TNF-α, IFN-γ, perforin and granzyme B (**Fig 3D**) suggesting that SPTBN2 expression in the TME may contribute to impaired TIL accumulation and function. Consistent with this hypothesis, we detected lesser levels of exhaustion markers such as LAG-3, TIGIT and PD-1 on the *Sptbn2*^-/-^ CD8+ T-cells from MC38 tumors (**Fig 3E**). Similar results were obtained in the immune-cold (MH6419c5) or immune-inflamed (MH6499c4) PDAC models in the subcutaneous or orthotopic settings (**Fig S3A-F**). Collectively, these results suggest that expression of SPTBN2 in the TME is associated with immunosuppression towards CTLs.

**Figure 3.**
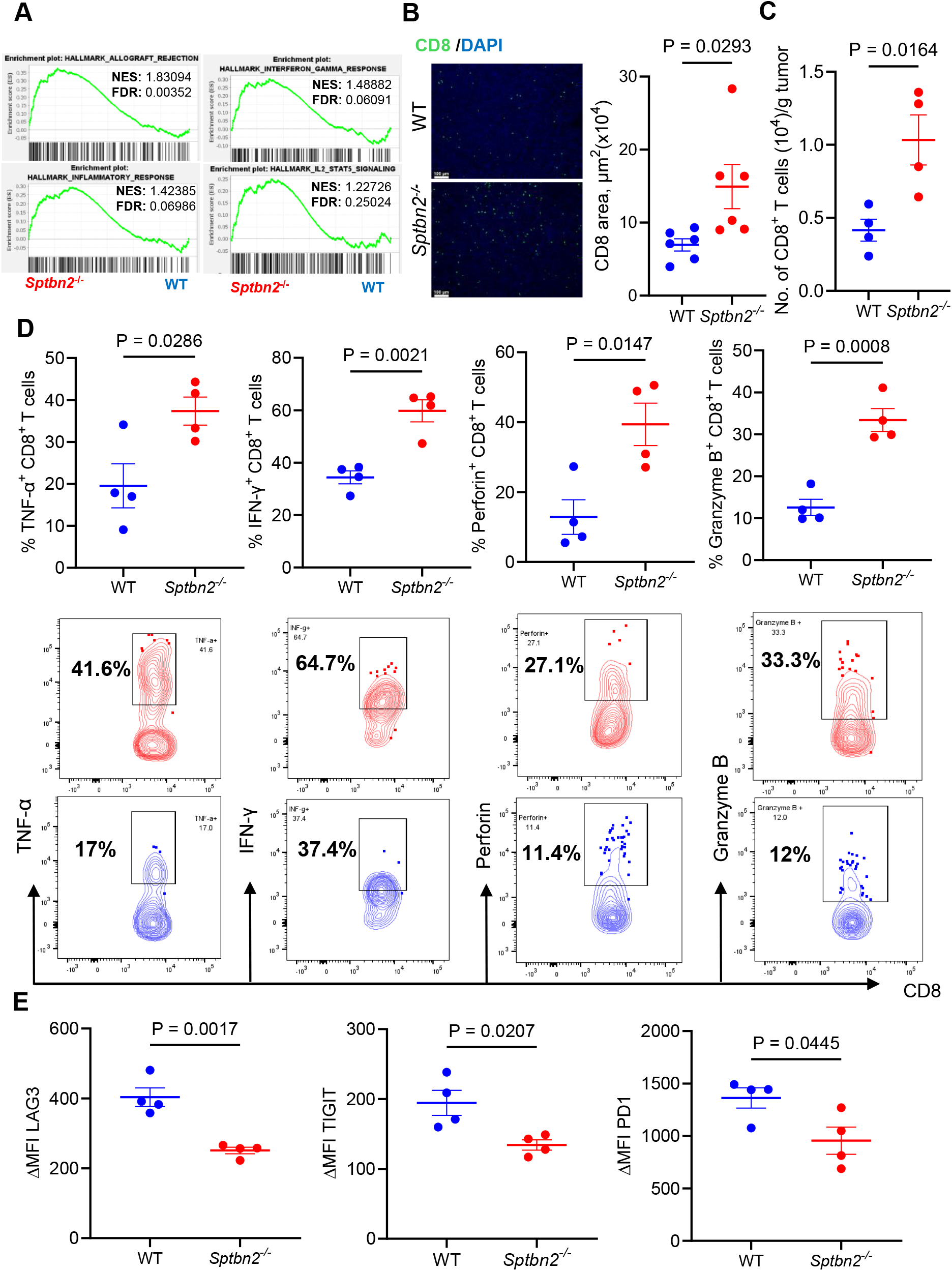
SPTBN2 inactivates tumor-infiltrating CTLs. A. GSEA of allograft rejection, interferon-γ response, inflammatory response, and IL2/STAT5 signaling hallmarks in MH6419c5 s.c. tumors grown in WT and *Sptbn2*^−^/^−^ mice. Normalized enrichment scores (NES) and false discovery rates (FDR) are shown. B. Representative immunofluorescence images of CD8 staining of MH6499c4 tumor growth in WT or *Sptbn2*^*-/-*^ mice (n=6), along with quantification of CD8-positive areas (Scale bar: 100µm). C. Flow cytometry analysis of CD8^+^ T-cell numbers per gram of the subcutaneous MC38 tumors grown in WT or *Sptbn2*^*-/-*^ mice (n=4). D. Flow cytometry analysis of the percentage of IFN-γ^+^, Perforin^+^, TNF-α^+^, Granzyme B^+^ of CD8^+^ T-cells isolated from subcutaneous MC38 tumor growth in WT or *Sptbn2*^*-/-*^ mice (n=4). E. Flow cytometry analysis of MFI of LAG3, TIGIT, and PD1 on CD8^+^ T-cells from subcutaneous MC38 tumors grown in WT or *Sptbn2*^*-/-*^ mice (n=4). Data are presented as mean± SEM. Statistical comparisons were conducted using the Student’s t-test.

Importantly, *Sptbn2*^-/-^ mice displayed notably decelerated growth of subcutaneous MC38 (**Fig 4A, S4A**) or MH6419c5 (**Fig 4B, S4B**) tumors compared to the wild type hosts. Similar results were observed in studies using MH6499c4 PDAC tumors in either subcutaneous (**Fig 4C, S4C**) or orthotopic (**Fig 4D**) settings. To determine whether effects of SPTBN2 expression were dependent on CD8+ T-cells, we depleted these cells from the host mice (**Fig 4E** and **Fig S4D**). Under these conditions, we observed a significant acceleration of tumor growth in *Sptbn2*^-/-^ mice (**Fig 4F-G**) suggesting that expression of SPTBN2 in the TME promotes tumor growth via inactivation of CD8+ CTLs.

**Figure 4.**
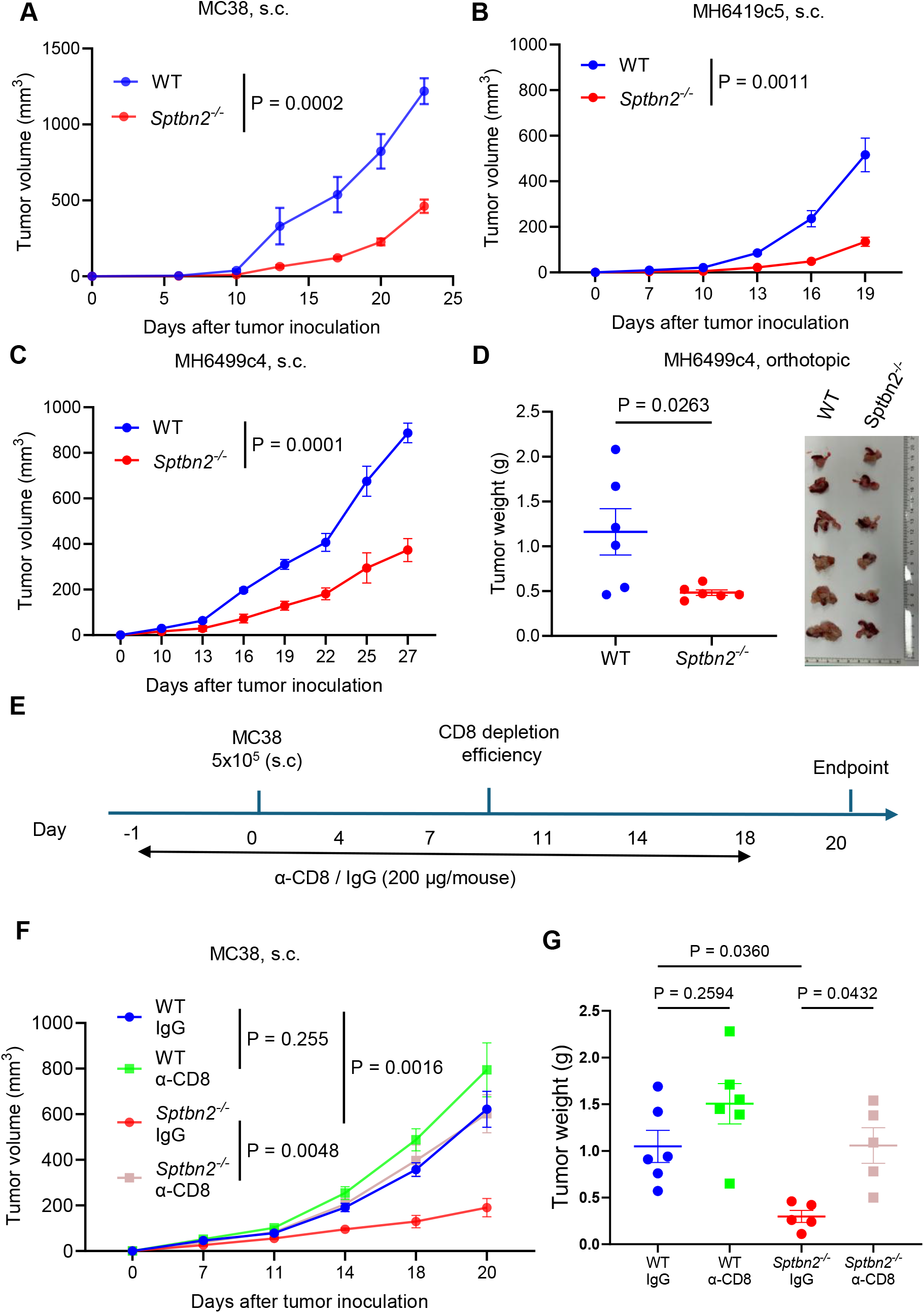
SPTBN2 promotes tumor growth in a CTL-dependent manner. A. Growth of subcutaneous MC38 tumors in WT and *Sptbn2*^*-/-*^ mice (n=4). B. Growth of subcutaneous MH6419c5 tumors in WT and *Sptbn2*^*-/-*^ mice (n=5). C. Growth of subcutaneous MH6499c4 tumors in WT and *Sptbn2*^*-/-*^ mice (n=4-5). D. Tumor weight of orthotopic MH6499c4 tumors in WT and *Sptbn2*^*-/-*^ mice (n=6). E. Schematic illustration of the treatment schedule for IgG (control) or α-CD8 antibodies in WT and. *Sptbn2*^*-/-*^ mice bearing MC38 subcutaneous tumors (n=5-6). F. Growth of subcutaneous MC38 tumors in WT and *Sptbn2*^*-/-*^ mice treated with either IgG or anti-CD8, as described in **E**. G. Tumor weight of subcutaneous MC38 tumors growth in WT and *Sptbn2*^*-/-*^ mice treated with either isotype control IgG or α-CD8, as described in **E**. Data are presented as mean± SEM. Statistical comparisons were conducted using the Student’s t-test for panels A-D and one-way ANOVA followed by Tukey’s post hoc test for panels F and G.

Anti-tumor CTLs play a pivotal role in ensuring the efficacy of diverse anti-cancer therapies including chemotherapy and immunotherapy ^3, 4, 22^. Given that expression of SPTBN2 is associated with CTL inactivation and tumor growth, we sought to determine whether SPTBN2 may also contribute to initial resistance to therapy. To this end, we used a “cold” MH6419c5 PDAC model, which was previously characterized to be insensitive to either chemotherapy with gemcitabine or immunotherapy with anti-PD1 antibody ^23, 24^. The latter treatment of the MH6419c5 tumor-bearing mice (**Fig 5A**) upregulated numbers of CD8+ TILs in both WT and *Sptbn2*^-/-^ mice (**Fig 5B**). Importantly, knockout of SPTBN2 in the TME resulted in increased levels of IFNγ-positive (**Fig 5C**) and Granzyme B-positive (**Fig 5D**) intratumoral CD8+ T-cells. Furthermore, these MH6419c5 PDAC tumors were resistant to immune checkpoint blockade in WT mice but responded with growth deceleration in *Sptbn2*^-/-^ mice (**Fig 5E, S5A-B**).

**Figure 5.**
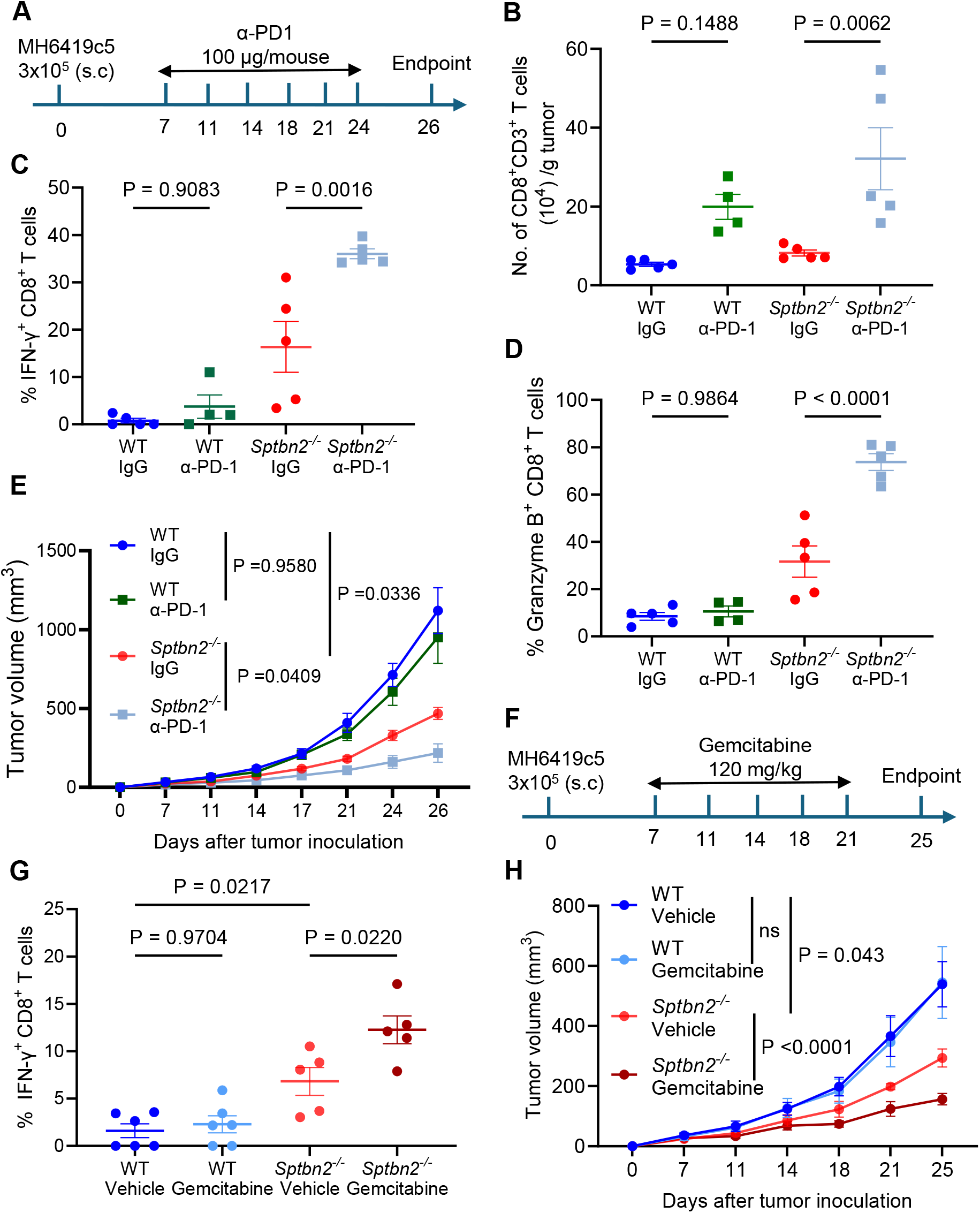
SPTBN2 confers PDAC resistance to chemotherapy and immune checkpoint blockade. A. Schematic illustration of the timeline for IgG or α-PD1 administration (100 μg/mouse, i.p.) in WT and *Sptbn2*^*-/-*^ mice bearing MH6419c5 subcutaneous tumors (n=4-5). B. Number of CD8^+^ T-cells per gram of subcutaneous MH6419c5 tumors in WT and *Sptbn2*^*-/-*^ mice treated with α-PD-1 or IgG, as described in panel **A**. C. Quantification of IFN-γ^+^ CD8^+^ T-cells in MH6419c5 subcutaneous tumors from WT and *Sptbn2*^*-/-*^ mice treated with α-PD-1 or IgG, as described in **A**. D. Quantification of Granzyme B^+^ CD8^+^ T-cells in MH6419c5 subcutaneous tumors from WT and *Sptbn2*^*-/-*^ mice treated with α-PD-1 or IgG, as described in **A**. E. Growth of subcutaneous MH6419c5 tumors in WT and *Sptbn2*^*-/-*^ mice treated with α-PD-1 or IgG, as described in **A**. F. Schematic diagram showing the timeline of Gemcitabine administration (120 mg/kg, i.p.) in WT and *Sptbn2*^*-/-*^ mice bearing MH6419c5 subcutaneous tumors (n=5-6). G. Analysis of IFN-γ^+^ cells within the CD8^+^ T-cell population in subcutaneous MH6419c5 tumors from WT and *Sptbn2*^*-/-*^ mice treated with gemcitabine or vehicle, as described in panel **F**. H. Growth of subcutaneous MH6419c5 tumors in WT and *Sptbn2*^*-/-*^ mice treated with Gemcitabine or vehicle, as described in **F**. Data are presented as mean± SEM. Statistical comparisons were conducted using one-way ANOVA followed by a Tukey post hoc test.

In the chemotherapy settings (**Fig 5F**), the course of five injections of gemcitabine affected the neither the total numbers of CD8+ TILs (**Fig S5C**) nor the frequency of IFNγ-expressing ones (**Fig. 5G**) in these tumors growing in WT mice. However, in line with data shown in **Fig 3**, knockout of SPTBN2 increased the number of intratumoral CD8+ T-cells and their expression of IFN-γ. Gemcitabine treatment further increased IFN-γ levels in the SPTBN2-deficient CTLs (**Fig 5G**). Importantly, whereas tumors in WT mice were resistant to gemcitabine, this treatment significantly inhibited tumor growth in *Sptbn2*^-/-^ mice (**Fig 5H-I, S5D**). These data suggest that expression of SPTBN2 in the TME impairs CTL activities in the context of chemotherapy and immunotherapy and contributes to the resistance to these therapeutic regimens.

### Inactivation of SPTBN2 in CAR T-cells maintains their memory state, supports their persistence and enhances therapeutic efficacy

Suppression of the intratumoral CTLs could occur indirectly through the effects of SPTBN2 expression in other cell types. To determine the importance of SPTBN2 expression specifically in the CTLs, we used splenic T-cells from WT or *Sptbn2*^-/-^ mice to generate anti-hCD19 CAR T-cells (**Fig S6A**). Compared to WT CAR T-cells, their *Sptbn2*-deficient counterparts expressed comparative levels of transgenic CAR (**Fig S6B**), modestly increased proliferation (**Fig S6C**) and increased expression of the central memory markers (**Fig 6A**). Importantly, upon co-culturing with hCD19-expressing target tumor cells, CAR T-cells derived from *Sptbn2*^-/-^ mouse responded with a greater production of IFN-γ and IL-2 cytokines (**Fig 6B**). Furthermore, under the co-culture conditions, WT CAR T-cells displayed significantly greater levels of LAG-3 and PD-1 exhaustion markers (**Fig 6C**) as well as lesser ability to kill target cells in a cytotoxic assay (**Fig 6D**) compared to *Sptbn2*^-/-^ CAR T-cells, suggesting that SPTBN2 inhibits CAR T-cell activity.

**Figure 6.**
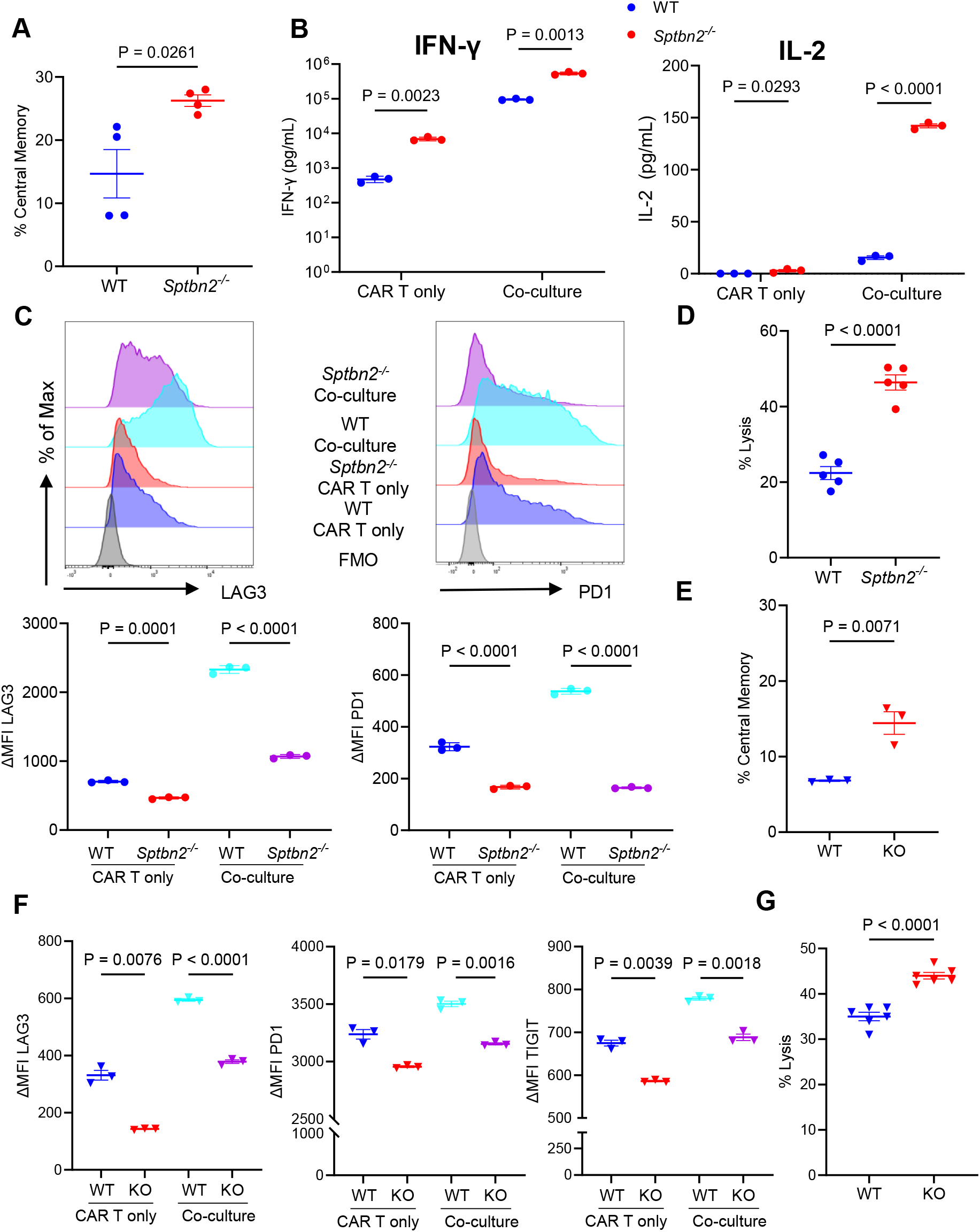
Expression of SPTBN2 in CAR T-cells promotes their inactivation and in vitro exhaustion. A. Quantification of central memory (CD62L^-^CD44^+^) markers in CD8^+^ CAR T-cells generated from WT and *Sptbn2*^*-/-*^ splenocytes (n=4). B. IFN-γ and IL-2 concentrations in the culture supernatants from the same co-culture experiment described in B (n=3). C. Representative flow cytometry analyses of MFI of LAG3 and PD-1 on CD8^+^ CAR T-cells cultured with or without B16F10.hCD19 tumor cells at an effector-to-target (E: T) ratio of 5:1 for 20 hours (n = 3). D. Cytotoxicity assay evaluating the killing efficiency of B16F10.hCD19-Luc target cells co-cultured with mouse WT or *Sptbn2*^*-/-*^ anti-CD19 CAR T-cells. Co-cultures were performed at an effector-to-target (E: T) ratio of 2:1 and assessed after 3 hours. E. Quantification of central memory (CD45RA^-^CCR7^+^) in CD8^+^ CAR T-cells generated from PBMC of healthy donors (n=3). F. Flow cytometry analyses of MFI of LAG3, PD-1, and TIGIT on CD8^+^ CAR T-cells cultured with or without OCI-Ly18 tumor cells at an effector-to-target (E: T) ratio of 5:1 for 20 hours (n = 3). G. Cytotoxicity assay evaluating the killing efficiency of human DLBCL OCI-Ly18-Luc target cells co-cultured with human WT or *SPTBN2* k.o. anti-CD19 CAR T-cells. Co-cultures were performed at an E: T ratio of 5:1 and assessed after 3 hours. Data are presented as mean± SEM. Statistical comparisons were conducted using Student’s t-test.

We next used experimental settings that did not involve the use of the *Sptbn2*^-/-^ mice. To this end, we generated human anti-hCD19 CAR T-cells and used a CRISPR approach to ablate SPTBN2 and generate *SPTBN2* k.o. CAR T cells (**Fig S6D-E**). Despite expressing similar levels of CAR (**Fig S6F**), there were greater numbers of central memory T-cells in *SPTBN2* k.o. CAR T-cells compared to WT CAR-T cells (**Fig 6E** and **S6G**). In addition, when co-cultured with target human diffuse large B-cell lymphoma (DLBCL) OCI-Ly18 cells that naturally express hCD19, the SPTBN2-deficient human CAR T-cells displayed fewer exhaustion markers (**Fig 6F** and **S6H**). Furthermore, *SPTBN2* k.o. CAR T-cells killed their target cells more efficiently than WT CAR T-cells (**Fig 6G**). These results indicate that SPTBN2 is an important negative regulator of human CAR T-cells.

The β-III spectrin protein encoded by *SPTBN2* regulates levels and localization of diverse cell surface proteins in different cell types ^25^. We sought to understand the mechanisms through which expression of SPTBN2 in CAR T-cells undermines their activities and promotes their exhaustion. First, we examined the involvement of SPTBN2 in effector trogocytosis – a process in which CAR T-cells acquire membrane patches containing tumor-associated antigens from target tumor cells ^26^. Spectrins were shown to promote cell membrane co-invasion and fusion in the context of Drosophila myoblast formation ^27^. These lipid membrane exchanges also play an important role in effector trogocytosis, which was previously characterized as a major mechanism for inactivating CAR T-cells ^28, 29^. Importantly, we observed that knockout of SPTBN2 in CAR T-cells restricted their ability to uptake CD19 antigen (**Fig 7A**) or lipid membrane fragments labeled with PKH67 dye (**Fig S7A**) from co-cultured B16F10-hCD19 target cells. These results indicate that SPTBN2 promotes effector trogocytosis.

**Figure 7.**
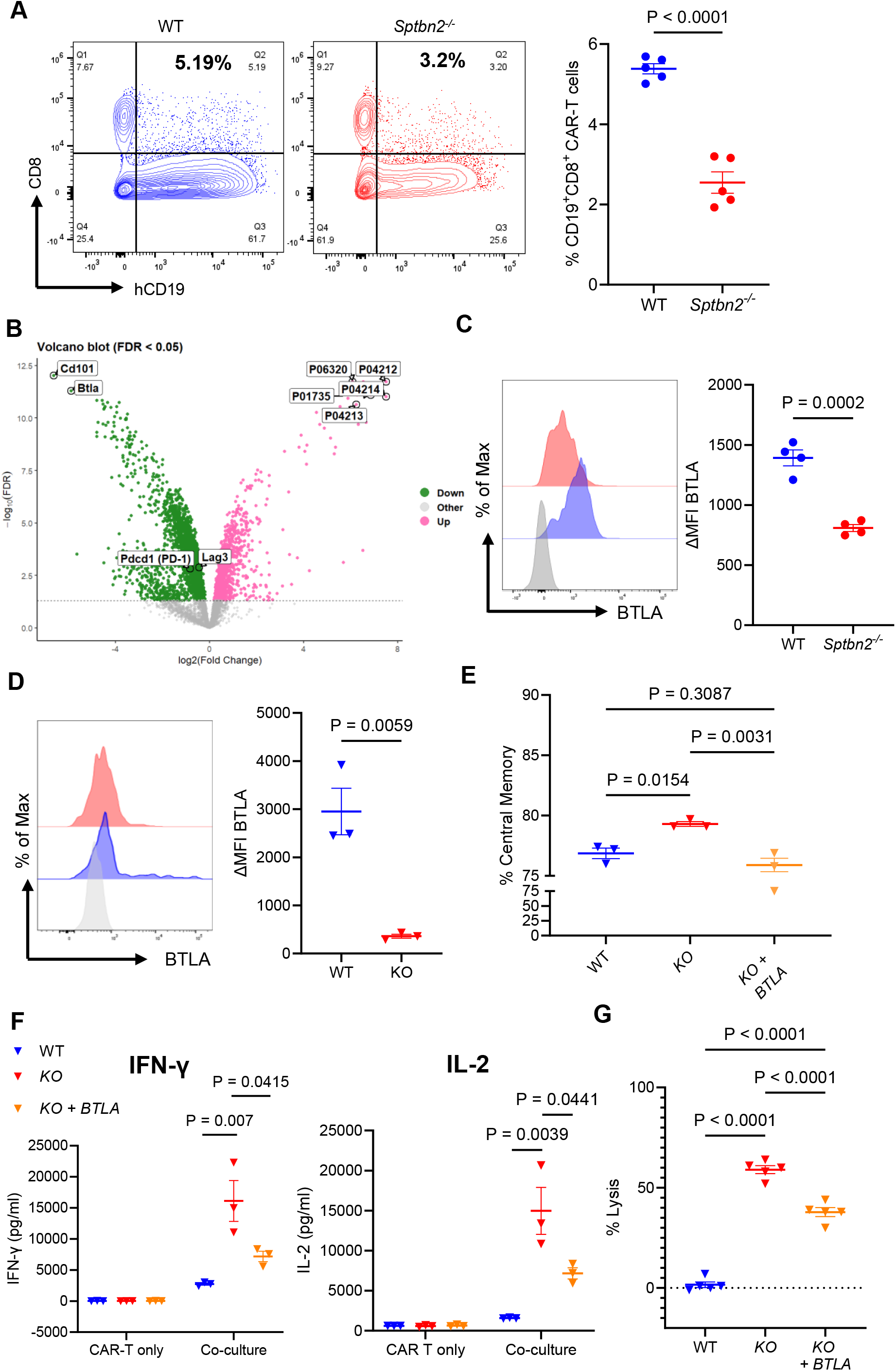
SPTBN2 regulates cell surface levels of plasma membrane proteins in CAR T-cells. A. Analysis of hCD19 transfer from B16F10.hCD19 tumor cells to CAR T-cells derived from WT or *Sptbn2*^*-/-*^ mice (n=5). CAR T-cells were co-cultured with B16F10.hCD19 cells at an E: T ratio of 5:1 for 15 minutes. The quantification represents the percentage of hCD19^+^ CD8^+^ CAR T-cells (n = 5). B. Volcano plot summarizing plasma membrane profiling results from CAR T-cells generated from WT or *Sptbn2*^*-/-*^ mice (n=5). C. Flow cytometry analysis of MFI of BTLA on mouse WT or *Sptbn2*^*-/-*^ CD8^+^ CAR T-cells D. Flow cytometry analysis of MFI of BTLA on human WT and SPTBN2 KO CD8^+^ CAR T-cells. E. Quantification of central memory (CD45RA^-^CCR7^+^) markers on indicated human CD8^+^ CAR T-cells (n=3). F. IL-2 and IFN-γ concentration in supernatants from indicated human CAR T-cells co-cultured with or without OCI-Ly18 cells for 24 h at an E: T ratio of 1:1 (n = 3) G. Cytotoxicity assay evaluating the killing of human DLBCL OCI-Ly18-Luc target cells co-cultured with indicated human WT or SPTBN2 KO anti-CD19 CAR T-cells. Co-cultures were performed at an effector-to-target (E: T) ratio of 1:1 and assessed after 18 hours. Data are presented as mean± SEM. Statistical comparisons were conducted using Student’s t-test for panels A, C, and D, and one-way ANOVA followed by Tukey’s post hoc test for panels E and G.

Second, we examined the cell surface protein repertoire regulated by SPTBN2 in CAR T-cells using a label-free plasma membrane profiling proteomics approach ^30, 31^ to compare the cell surface levels of proteins in WT and *Sptbn2*^-/-^ CAR T-cells (**Fig 7B**). Quantitative mass spectrometry analysis revealed that knockout of SPTBN2 affected the levels of numerous plasma membrane proteins (**Fig S7B-C, Supplemental Table 2**). Several species of T-cell receptor beta chain were notably increased in *Sptbn2*^-/-^ CAR T-cells, whereas these cells also displayed a significant decrease in numerous cell surface proteins including BTLA and CD101 (**Fig 7B**). CD101 acts as an inhibitor of T-cell function via suppressing production of IL-2 ^32^; this characteristic is in line with increased IL-2 production by SPTBN2-deficient CAR T-cells (**Fig 6B**).

We decided to focus on B and T lymphocyte attenuator (BTLA) because it is a key negative regulator of T lymphocyte activation ^33, 34^ and of CAR T-cell activities ^35^. Flow cytometry analysis confirmed that the cell surface of BTLA was decreased upon SPTBN2 ablation in murine (**Fig 7C**) and human (**Fig 7D**) CAR T-cells. Importantly, re-expression of BTLA in the SPTBN2-deficient human CAR T-cells (**Fig S7D**) abrogated an increase in the central memory state markers (**Fig 7E** and **S7E**). Furthermore, production of cytokines (**Fig 7F**) and ability to kill target DLBCL OCI-Ly18 cells (**Fig 7G**) were significantly attenuated in the BTLA-expressing *SPTBN2* k.o. CAR T-cells. These results suggest that SPTBN2-dependent expression of BTLA on the CAR T-cells is at least in part responsible for their inactivation.

Success of clinical treatments using CAR T-cells is linked to their expansion and persistence, which is in turn associated with memory states of these cells ^36, 37^. We examined the association between SPTBN2 expression in manufactured anti-CD19 CAR T cells with their persistence and efficacy (manifested by B cell aplasia, BCA) in pediatric patients with acute lymphocytic leukemia ^38^. At the single-cell level, baseline (**Fig 8A**) or specific antigen-restimulated (**Fig S8A**) manufactured CAR T cells from patients with shorter BCA durations exhibited significantly higher *SPTBN2* expression compared to those from patients with long-term BCA (≥5 years). This trend was also observed at the patient level (**Fig S8B**). Together, these findings suggest that elevated *SPTBN2* expression is associated with reduced persistence and clinical failure of CAR T cells.

**Figure 8.**
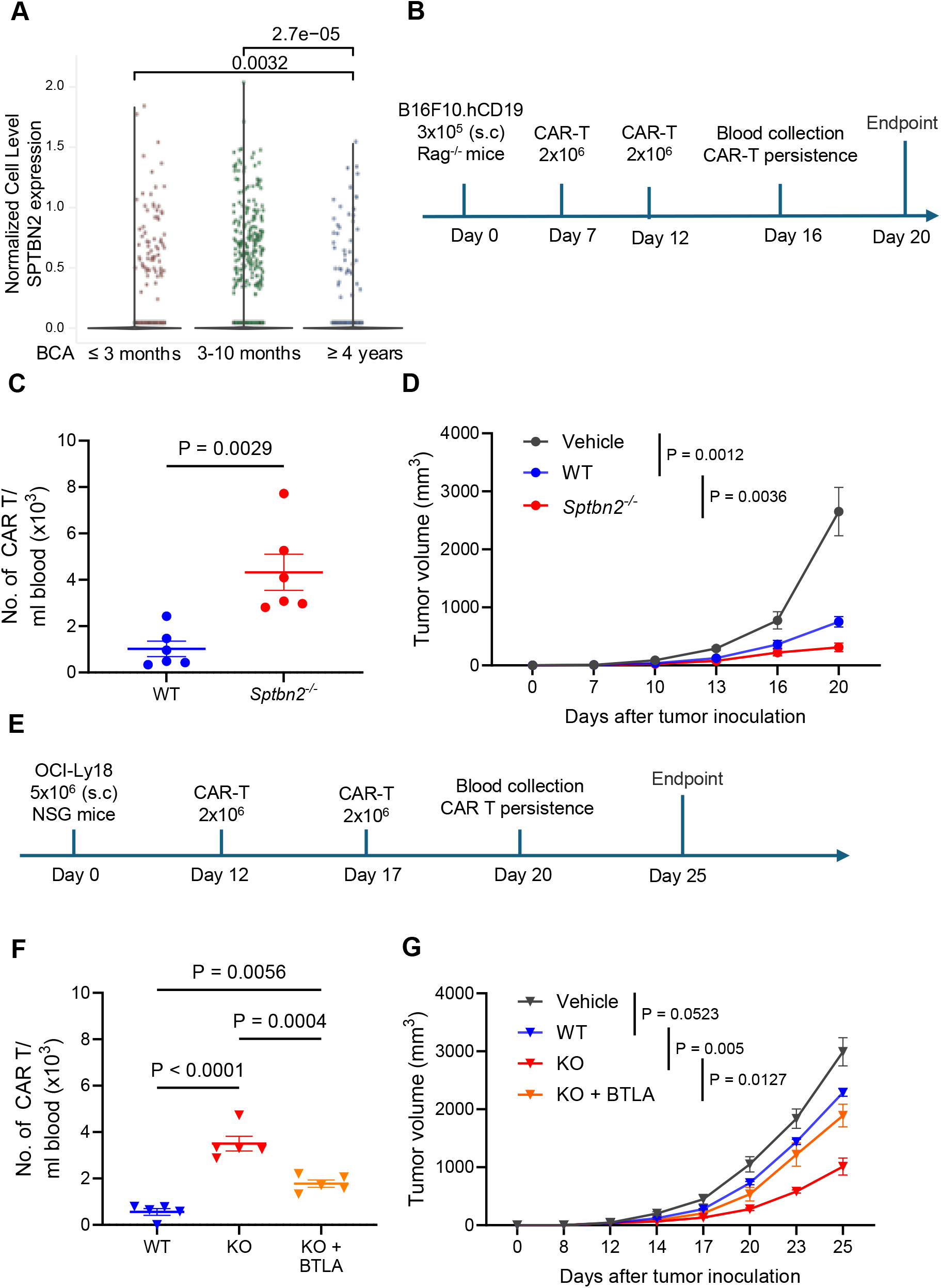
Inactivation of SPTBN2 in CAR T-cells supports their persistence and enhances therapeutic efficacy. A. Violin plot of normalized SPTBN2 expression in manufactured CAR T cells stratified by B-cell Aplasia (BCA) outcome group defined in ^38^ (GSE262072). Each dot represents a cell. Expression of SPTBN2 in manufactured CAR T cells cultured in the basal unstimulated condition are shown. P-values shown are unadjusted. B. Schematic illustration of the CAR T-cell treatment schedule in Rag^-/-^ mice bearing B16F10.hCD19 tumors (n=6). C. Quantification of CAR T/ml blood of tumor-bearing mice treated with WT or Sptbn2^-/-^ CAR T-cells collected on day 16, as described in **A**. D. Tumor growth curves showing the volume of B16F10.hCD19 tumors in Rag1^-/-^ mice treated with WT or *Sptbn2*^-/-^ CAR T-cells, following the treatment schedule described in **A**. E. Schematic illustration of the CAR T-cell treatment schedule in NSG mice bearing human DLBCL OCI-Ly18 tumors (n=5). F. Quantification of CAR T/ml blood of tumor-bearing mice treated with WT or SPTBN2 KO CAR T-cells collected on day 20, as described in **D**. G. Tumor growth curves showing the volume of OCI-Ly18 tumors in NSG mice treated with WT or SPTBN2 KO CAR T-cells, following the treatment schedule described in **D**. Data are presented as mean± SEM. Statistical comparisons were conducted using a two-sided unpaired Wilcoxon rank sum test for panel A, Student’s t-test for panels B-C, and one-way ANOVA followed by Tukey’s post hoc test for panels E-F.

Further *in vivo* studies were conducted to reveal the importance of SPTBN2 in the persistence and efficacy of CAR T-cell anti-tumor treatments. We first treated the B16F10-hCD19 tumor-bearing *Rag1*^-/-^ mice with mouse anti-hCD19 CAR T-cells generated from either WT or *Sptbn2*^-/-^ splenic T-cells on days 7 and 12 after tumor inoculation (**Fig 8B**). Analysis of peripheral blood collected on Day 16 revealed a greater number of *Sptbn2*^-/-^ CAR T-cells compared to WT ones (**Fig 8C**). Furthermore, analyses of splenic (**Fig S8C**) and tumor (**Fig S8D**) tissues harvested on Day 20 also revealed a greater persistence of the *Sptbn2*-deficient CAR T-cells. Importantly, adoptive transfer of these cells significantly decelerated tumor growth compared to either vehicle treatment or infusion of WT CAR T-cells (**Fig 8D** and **S8E**).

In the second model, NOD scid gamma (NSG) mice were subcutaneously inoculated with human DLBCL OCI-Ly18 cells and then 12 and 17 days later received the adoptive transfer with WT or SPTBN2 k.o. human CAR T-cells (**Fig 8E**). Under these conditions, knockout of SPTBN2 significantly increased the presence of CAR T-cells (**Fig 8F** and **S8F**) and their infiltration of tumors (**Fig S8G**) and improved their ability to control tumor growth (**Fig 8G** and **S8H**). Importantly, re-expression of BTLA reversed these phenotypes (**Fig 8F-G** and **S8F-G-H**). Collectively, these results suggest that SPTBN2 inactivates CAR T-cells through the mechanisms that involve BTLA and that targeting SPTBN2 in the context of the CAR T-cell adoptive transfer therapy increases its efficacy.

## DISCUSSION

A totality of conflicting tumor-promoting and tumor-suppressing functions of cell surface proteins ultimately determines the course for tumor development, growth and progression. In principle, as a global organizer of interaction between signaling receptors and actin and other cytoskeletal elements of, SPTBN2 upregulation would be expected to enhance the receptor-driven signal transduction in both pro-tumorigenic and anti-tumorigenic pathways. However, growing tumors appear to exploit additional selective mechanisms and utilize the SPTBN2-driven changes in malignant cells to stimulate their proliferation and survival ^16-19^.

Likewise, as demonstrated in this study, for the cells of the TME such as intratumoral CTLs, upregulation of SPTBN2 drives their exhaustion and inactivation leading to immunosuppression and stimulation of tumor growth and resistance to anti-cancer therapies. Furthermore, expression of SPTBN2 in the TME is important for efficient tumor angiogenesis and stromagenesis. These results characterize SPTBN2 as a pivotal regulator of tumor-induced immunosuppression and therapeutic resistance.

Our data suggest that induction of SPTBN2 stimulated by nutritional deficit in activated intratumoral CTLs acts to limit their anti-tumor activity and push them towards exhaustion. Outside of the tumor context, the biological significance of this regulation remains to be understood. Perhaps upregulation of SPTBN2 under conditions of impaired perfusion (for example, in inflamed tissue) that leads to nutrient deprivation may result in a prevailing stabilization of cell surface proteins that possess immune-regulatory function. Such proteins (e.g. BTLA) can perhaps act to safeguard against excessive immunopathology and tissue damage in the foci of inflammation. While, unlike BTLA knockout ^39^, no inflammatory phenotypes were observed in unchallenged *Sptbn2*-null mice, this might reflect the involvement of additional anti-inflammatory proteins whose cell surface levels are regulated by SPTBN2. Furthermore, although SPTBN2 is not essential to prevent inflammatory phenotypes in naïve mice, it would be of interest to challenge these *Sptbn2*^-/-^ mice with a range of inflammatory and infectious stimuli that might reveal additional important functions of this gene. Furthermore, whereas current observations suggest a role for BTLA, future mechanistic studies focusing on other potential mediators of SPTBN2 induction such as CD101 and others are warranted.

Current studies in solid tumors such as colon and pancreatic ductal adenocarcinomas demonstrate that the induction of SPTBN2 leading to inactivation of the tumor-infiltrating CTLs represents a previously unrecognized mechanism of immunosuppression in TME. In human tumors, levels of SPTBN2 inversely correlate with T-cell infiltration manifested by expression of *CD8A*. In mouse models, native tumor-infiltrating CD8+ T-cells and CAR-bearing T-cells that express SPTBN2 display increased markers of exhaustion and impaired production of cytokines and cytotoxic markers. Accordingly, expression of SPTBN2 in the TME cells notably accelerates tumor growth. Furthermore, given that CTLs are essential for robust responses to chemotherapy and immune checkpoint inhibitors ^3, 4, 22^, it is not surprising to observe that knockout of SPTBN2 reverses cross-resistance of pancreatic tumors to these treatment regimens. These observations argue for development of therapeutics to inhibit SPTBN2 function.

This work has several ***limitations*** that will be addressed in future studies. First, the mechanistic underpinnings of *SPTBN2* upregulation in the malignant and non-malignant cells within tumor are yet to be fully determined. In cancer cells (e.g. pancreatic cancer) it has been suggested that de-methylation of the SPTBN2 promoter may be involved ^19^. *SPTBN2* was among the genes that were upregulated in the endothelial cells upon exposure to the tumor-derived extracellular vesicles ^40^. Here we demonstrate that conditions of nutritional deficit induce *SPTBN2* expression in CD8+ T-cells. However, the precise mechanisms by which the non-malignant cells of the TME including CTLs upregulate *SPTBN2* remain to be delineated. Identification and characterization of such mechanisms may help to target their critical regulators to prevent *SPTBN2* induction, subsequent stimulation of immunosuppression, angiogenesis, stromagenesis and resulting ensuing tumor growth. Likewise, better understanding of how SPTBN2 in the intratumoral endothelial cells and CAFs can promote angiogenesis and stromagenesis warrants future studies.

Second, the molecular basis by which SPTBN2, classically a cortical cytoskeletal scaffold, regulates membrane protein stability, trafficking, or signaling in CD8^+^ T cells remain to be fully elucidated. Lastly, additional studies are also required for delineating the mechanisms by which SPTBN2 expression destabilizes the effector memory phenotypes. It is plausible that SPTBN2-mediated cell surface stabilization of receptors capable of initiating the phosphoinositide 3-kinase-AKT signaling cascade may negatively affect transcription driven by the AKT substrate FOXO1, as these pathways are essential for regulating memory programs in T cells ^37^.

Data presented here also highlights the importance of SPTBN2 in undermining the efficacy of adoptive transfer therapy using CAR T-cells. Limited persistence and terminal exhaustion of CAR T-cells *in vivo* associated with the loss of T-cell memory gene signatures represent a major roadblock for successful adoptive cell therapy using these cells ^37, 41, 42^. Our studies reveal that SPTBN2 acts as an important regulator of memory programming and exhaustion in CAR T-cells. Furthermore, SPTBN2 expression in manufactured CAR T cells is associated with their clinical failure. Accordingly, expression of SPTBN2 in CAR T cells undermines their anti-tumor activity. Conversely, gene editing-based inactivation of SPTBN2 in CAR T cells is associated with a favorable memory profile, increases persistence, and ultimately improves the anti-tumor activity CAR T cell therapy in multiple models. These data provide impetus to further explore genetic or pharmacologic approaches for inactivation of SPTBN2 *ex vivo* in CAR T cells prior to infusion with the goal of improving treatment outcomes in cancer patients.

## ACKNOWLEDGEMENTS

This work was supported by NIH/NCI grants R01 CA285321 (to S.Y.F.) and R01 CA288849 (to V.S.S. and S.Y.F.). We thank Dr. Andy Minn (University of Pennsylvania) for reagents and the members of Fuchs, Koumenis, Stanger, Guo, Conn, Atherton, and Fan labs for critical discussion.

## MATERIAL AND METHODS

### Study approvals

Use of human T-cells that were previously collected from healthy donors under informed consent and could not be directly or indirectly linked to individual human participants was approved for the Human Immunology Core by the Institutional Review Board of the University of Pennsylvania. All animal experiments were approved by the Institutional Animal Care and Use Committee of the University of Pennsylvania (protocol #803995) and were carried out in accordance with the IACUC guidelines.

#### Cell culture

Mouse colon adenocarcinoma MC38 cell line was purchased from Kearfast and maintained according to the recommendations. Mouse MH6140c5 and MH6499c4 PDAC cell lines were previously described ^24^. hCD19-B16F10 cells were engineered to express Firefly luciferase as described previously ^43^. All those cells were cultured in DMEM (Gibco, #11965-092), supplemented with 10% HyClone fetal bovine serum (Cytiva, #SH30071.03), 1× Antibiotic/Antimycotic Solution (Cytiva, #SV3007901). OCI-Ly18/Luciferase provided by Dr. Marco Ruella (Perelman Center for Advanced Medicine, University of Pennsylvania, PA, USA) was maintained in RPMI-1640 Medium (Gibco, #22400089), supplemented with 10% HyClone fetal bovine serum (Cytiva, #SH30071.03), 1mM sodium pyruvate (Gibco, #11360070),1× Antibiotic/Antimycotic Solution (Cytiva, #SV3007901).

Mouse T-cells were cultured in RPMI-1640 Medium (Gibco, #22400089), supplemented with 10% HyClone fetal bovine serum (Cytiva, #SH30071.03), 1mM sodium pyruvate (Gibco, #11360070),1× Antibiotic/Antimycotic Solution(Cytiva, #SV3007901), 55μM 2-mercaptoethanol (Sigma, #60242), and 20ng/mL recombinant mouse IL-2 (Biolegend, #575408). Mouse endothelial cells were cultured in advanced DMEM with reduced serum (Gibco, #12491015), supplemented with 15% HyClone fetal bovine serum (Cytiva, #SH30071.03), 2mM L-Glutamine (Gibco #25030081), 1× Antibiotic/Antimycotic Solution (Cytiva, #SV3007901), 0.1 mg/ml endothelial cell growth factor (Merck, #02-102), and 1 IU/ml heparin (Sigma, #H5515-25KU). Human T-cells obtained from the healthy donors by the Human Immunology Core of the University of Pennsylvania were cultured in RPMI-1640 Medium (Gibco, #22400089), supplemented with 10% HyClone fetal bovine serum (Cytiva, #SH30071.03), 1mM sodium pyruvate (Gibco, #11360070),1× Antibiotic/Antimycotic Solution(Cytiva, #SV3007901), 55μM 2-mercaptoethanol (Sigma, #60242), 5 ng/mL recombinant human IL-7 (PeproTech, #200-07-50UG), and 5 ng/mL recombinant human IL-15 (PeproTech, #200-15-50UG).

### Animals

C57BL/6J, *Rag1*^*-/-*^, *Sptbn2*^-/-^ (C57BL/6 genetic background), and NSG mice were obtained from the Jackson Laboratory. All animal experiments were conducted using both male and female mice between 8 and 10 weeks of age. Mice were housed under specific pathogen-free (SPF) conditions with a 12-hour light/dark cycle and maintained at a temperature of 20–25°C. They had ad libitum access to water and standard laboratory chow (LabDiet 5010 and MF diet; Animal Specialties & Provisions), in accordance with the guidelines of the American Association for Laboratory Animal Science (AALAS). Littermates from different cages were randomly assigned to experimental groups and were housed under identical environmental conditions. Animal health and welfare were routinely monitored by qualified veterinary staff.

### Isolation, culture, and tube formation analyses of primary lung ECs

The ECs were carried out similarly to those previously described ^40^. Briefly, lungs were collected from 3-week-old wild-type (WT) and *Sptbn2*^*-/-*^ mice, washed with ice-cold SFM medium, and dissected into small pieces using dissection scissors. The tissue was incubated in a digestion solution containing 2 mg/ml Collagenase II (MP Biomedicals, #100502) and 100 µg/ml DNase I (Roche, #10104159001) for 40 minutes with continuous agitation at 200 RPM at 37°C. Digestion was stopped by adding prewarmed FBS, followed by passing through 70-μm filters and centrifugation at 500 ×g for 5 minutes. The cell pellet was resuspended in 5 mL cold PBS and washed twice. The cells were seeded into a 150mm dish and incubated overnight. Non-attached cells were removed, and the remaining cells were washed with PBS and cultured in ECs media. Upon confluence, the cells were collected using a non-enzyme detachment solution and enriched using CD31 Microbeads (Miltenyi Biotec, #130-097-418). After confirming the purity, the enriched cells were expanded in ECs media for experiments.

For tube formation, ECs were washed twice with PBS and seeded into 24-well plates (8 × 10^4^ cells per well) coated with Cultrex RGF BME, Type 2 (R&D, #3533-005-02). Images of tube formation were captured 6 hours after seeding using contrast phase microscopy, and the results were analyzed by ImageJ software

### Nutrient stress in mouse and human CD8^+^ T-cells

The cells were cultured under the control condition or deficient conditions (arginine, branched chain amino acid, glucose, glutamine, or methionine deprivation) for 3 hours as reported ^7^. For polysome profiling via total RNA-seq of nutrient stress conditions, RNA samples were sequenced at the Center for Applied Genomics at the Children’s Hospital of Philadelphia. These samples were then used for RNA sequencing, carried out as previously described ^7^.

### Mouse and human CAR T-cell preparation and *in vitro* experiments

Mouse CAR T-cells were prepared similarly to those described previously ^44^. Briefly, the splenocytes were collected from WT or *Sptbn2*^*-/-*^ mice. Isolated T-cells were stimulated with Dynabeads Mouse T-Activator CD3/CD28 beads (GIBCO, cat# 11453D) for 48 hours and were separated from the beads before transduction. The adenovirus (hCD19-BBz) was subsequently prepared on Phoenix cells and added to retronectin-precoated (20 µg/mL, Takara, #T100B) 24-well plates. After being spun down at 2000 ×g for 1 hour, the T-cells were added for transduction and expanded for three days before being harvested. Transduction efficiency was validated through flow cytometry analysis. The validated CAR T-cells, 3 to 8 days post-transduction, were used for both in vitro and in vivo experiments with the B16F10.hCD19 tumor model.

For human CAR T, CD8^+^ and CD4^+^ T-cells were isolated from healthy donors, provided by the Human Immunology Core (University of Pennsylvania, PA, USA). The cells were mixed at an appropriate ratio and cultured in full human T-cell culture media (TCM). The following day, 1×10^7^ T-cells were collected and washed twice with reduced-serum minimal essential medium (Opti-MEM, Life Technologies, #31985062) before being electroporated with the ribonucleoprotein (RNP) complex, consisting of pre-assembled SPTBN2-targeting sgRNA and Cas9 nuclease (IDT), along with an electroporation enhancer (IDT). Electroporation was performed using the P3 Primary Cell 4D-Nucleofector X Kit (Lonza, #V4XP-3024). Cells were immediately transferred to 1 mL of pre-warmed complete medium for recovery and incubated for 48 hours. Then, T-cells were stimulated using Dynabeads Human T-Activator CD3/CD28 (Invitrogen, #11132D) at a 1:3 T-cell-to-bead ratio for 24 hours, followed by transduction with a CAR.hCD19 or CAR.hCD19-P2A-BTLA-encoding lentivirus. The beads were subsequently removed, and the cells were harvested for quality assessment. Transduction efficiency was quantified by flow cytometry. Cell pellets were collected for Western blot and qPCR analyses to verify KO efficiency. Verified cells were then expanded in TCM for subsequent experiments with the OCI-Ly18 tumor model.

**Cytotoxicity assays** were carried out as previously described ^43, 44^. Briefly, human or mouse CAR T-cells were pre-treated with adenosine (1mM, 24hr) to mimic the immunosuppressive conditions of the TME. After that, CAR T-cells were washed and co-cultured with pre-seeded Luciferase-expressing tumor cells in 96-well plates with a total volume of 200 μL at a ratio as indicated. Target cells alone were seeded in parallel at the same density to quantify spontaneous death luciferase expression (measured in relative luminescent units; spontaneous death RLU). Additionally, target cells with water were included as a control for maximal killing (maximal killing RLU). Following coculture, 100 μl of luciferase substrate (Promega, #E6120) was added to the remaining supernatant and cells. Luminescence was measured after a 10-minute incubation. Percent cell lysis was calculated using the formula: % lysis = 100 × (spontaneous death RLU - test RLU) / (spontaneous death RLU - maximal killing RLU).

**For cytokine secretion assays**, pre-seeded tumor cells were co-cultured with CAR T-cells at the indicated effector-to-target ratios. Target cells cultured alone under the same conditions were included as controls. After 20-24 hours of co-culture, supernatants were collected for ELISA to quantify cytokine secretion. For mouse CAR T-cells, IL-2 and IFN-γ concentrations were measured using the ELISA MAX™ Standard Set Mouse IL-2 (BioLegend, #431001) and ELISA MAX™ Standard Set Mouse IFN-γ (BioLegend, #430801). For human CAR T-cells, IL-2 and IFN-γ concentrations were measured using the ELISA MAX™ Standard Set Human IL-2 (BioLegend, #431801) and ELISA MAX™ Standard Set Human IFN-γ (BioLegend, #430101). All ELISA analyses were performed in accordance with the manufacturer’s instructions.

**For exhaustion analysis**, pre-seeded tumor cells were co-cultured with CAR T-cells at the indicated effector-to-target ratios. After 20–24 hours, cells were collected, washed twice with PBS, stained with an antibody cocktail, and analyzed by flow cytometry.

**For trogocytosis**, CAR T-cells were co-cultured with target cells at a 1:5 ratio for 15 minutes. After co-culture, cells were washed with PBS, stained with Fixable viability dyes (FVD), followed by CD8 and hCD19 staining. After staining, cells were washed again with PBS and analyzed by flow cytometry. Trogocytosis was assessed by evaluating the acquisition of hCD19 antigens by CAR T-cells, as well as the loss of CD19 expression on target tumor cells, indicating the transfer of membrane-bound antigen. Similarly, for membrane transfer analysis, PKH67-labeled tumor cells were co-cultured with CAR T-cells at a 5:1 ratio for 15 minutes. Cells were then collected, and membrane transfer was quantified by measuring PKH67 acquisition by CAR T-cells.

### Plasma membrane profiling proteomic screen

WT and *Sptbn2*^-/-^ CAR T-cells were collected, and dead cells were removed using the Dead Cell Removal Kit (Miltenyi Biotec, #130-090-101) according to the manufacturer’s instructions. Cells were then washed three times with ice-cold PBS. Plasma membrane proteins were labeled and isolated using the Pierce™ Cell Surface Protein Biotinylation and Isolation Kit (Thermo Fisher Scientific, #A44390) following the manufacturer’s protocol.

#### Protein Digestion

Proteins were eluted using lysis buffer containing 1% sodium dodecyl sulfate (SDS, Affymetrix), 100mM Tris-HCl (Invitrogen), and 10 mM Dithiothreitol with room temperature incubation for 45 minutes. Columns were centrifuged at 2000 g for 2 minutes to elute proteins. SDS concentration was adjusted to 3%. Protein concentration was estimated with an SDS-Page gel stained with Bradford Coomassie solution. Intensity analysis was performed using GelAnalyzer 19.1 against a serial dilution of an in-house generated E. coli lysate standard. Proteins were processed using the S-Trap protocol ^45^, including reduction with TCEP, alkylation with iodoacetamide, and acidification prior to loading onto S-Trap columns. Samples were digested overnight at 37°C with trypsin and LysC. Peptides were then eluted, dried, desalted using C18 StageTips (T3, Affinisep), and reconstituted in 0.1% TFA containing iRT peptides. Peptide concentration was determined by OD280 on a Synergy H1 microplate reader (BioTek), and samples were normalized for injection.

#### Mass Spectrometry Data Acquisition

Samples were randomized and analyzed on a timsTOF HT mass spectrometer coupled to a nanoElute 2 system. Peptides were separated on a 25 cm C18 column using a 35 min gradient at 300 nL/min. Data were acquired in DIA-PASEF mode over an m/z range of 100–1700 and mobility range of 0.70–1.30 1/K_0_, with a cycle time of 0.65 s. Collision energy was applied as a mobility-dependent ramp. The ion mobility dimension was calibrated using Agilent ESI tuning mix standards.

#### Quality Control and Data Processing

The suitability of the instrumentation was monitored using QuiC software (Biognosys; Schlieren, Switzerland) for the analysis of the spiked-in iRT peptides. As a measure for quality control, standard human K562 digest was injected between samples and data was collected in data dependent acquisition (DIA) mode. The data collected were analyzed in Bruker Proteoscape to track the quality of the instrumentation. MS/MS raw files were processed in Spectronaut v.20.1 (Biognosys AG) using direct DIA mode ^46^. A reference mouse proteome including 25,508 canonical proteins and reviewed isoforms from UniProt was used, appended with the list of 245 common protein contaminants. Trypsin was specified as an enzyme with two possible missed cleavages. Carbamidomethyl of cysteine was specified as fixed modification and protein N-terminal acetylation and oxidation of methionine were considered variable modifications. The false discovery rate limit of 1% was set for peptide and protein identification. The rest of the search parameters were set to the default values.

#### Bioinformatics Analysis

Proteomics data processing and statistical analysis were conducted in R. The MS2 intensity values generated by Spectronaut were utilized for analyzing the entire proteome dataset. The data were log2-transformed and normalized by subtracting the median value for each sample. To ensure data quality, proteins were filtered to retain those with complete detection in at least one cohort. Differential protein abundance across groups was assessed using a Limma t-test, and the results were visualized using volcano plots. Lists of differentially abundant proteins were generated based on criteria of adjusted P.Value <0.05, resulting in a prioritized list for subsequent bioinformatics analysis. All relevant mass spectrometry proteomics data have been deposited to the ProteomeXchange Consortium via the PRIDE partner repository with the dataset identifier ***PXD076271***.

### RNA-seq

Total RNA was extracted with the RNeasy Plus Mini Kit (QIAGEN, 74134), following the manufacturer’s instructions. RNA concentration and purity were assessed using a Nanodrop spectrophotometer. RNA-seq libraries were prepared using the Illumina Total RNA Stranded Ligation with Ribo-Zero^+^ kit (Illumina, #20040529). The prepared libraries were pooled and sequenced on an Illumina NextSeq 2000 system using a P2 100-cycle flow cell in a single-end mode. Raw sequencing reads were subjected to quality control using FastQC v0.12.1. Adapter sequences and low-quality bases were trimmed using Trim Galore or Cutadapt v4.0. Transcript abundances were quantified with kallisto v0.51.1, referencing the corresponding *Musculus* reference genome (GRCm39). Transcript-to-gene counts were imported into R using tximport v1.36.1, and lowly expressed genes (counts per million > 1 at least 3 samples) were filtered before normalization and differential expression analysis with edgeR v4.6.3. Genes with an adjusted p-value < 0.05 (Benjamini–Hochberg correction) and an absolute log2 fold-change ≥ 1 were considered significantly differentially expressed. Gene Ontology (GO) and pathway enrichment analyses were performed using the clusterProfiler R package. Heatmaps and volcano plots were generated using ggplot2 and pheatmap to visualize gene expression patterns. GSEA v4.4.0 was used to perform Hallmark gene set enrichment analysis. All relevant data were deposited to the GEO repository under accession number ***GSE326322***.

### Tumor growth and therapy studies

For syngeneic s.c. tumor model, 5×10^5^ MC38, 3×10^5^ MH6499c4, 3×10^5^ MH6149c5 were subcutaneously inoculated into the right flank of C57BL/6J mice. Tumor volume and body weight changes were estimated as 0.5 × length × width^2^, twice a week, until the endpoint of the experiments.

For the orthotopic tumor model, experimental mice were anesthetized with isoflurane inhalation under sterile surgical conditions. A small incision was made in the left abdominal flank to expose the tail of the pancreas. A suspension of 3 × 10^5^ MH6499c4 cells was slowly injected into the pancreatic tail using a 30-gauge needle. Following successful implantation, the pancreas was gently repositioned into the peritoneal cavity, and the abdominal wall and skin were carefully sutured. Mice were monitored twice weekly for signs of distress and loss of body weight. Tumor masses were measured at the endpoint of the experiment.

For the CAR T-cell therapy experiments, 3 × 10^5^ B16F10.hCD19 cells were subcutaneously injected into the right flank of *Rag1*^*-/-*^ mice. CAR T-cells were generated from splenocytes isolated from either WT or *Sptbn2*^*-/-*^ mice. Two intravenous doses of CAR T-cells were administered to tumor-bearing mice on day 7 and day 15 post-tumor implantation. When tumors reached approximately 1500 mm^3^, mice were euthanized, and tumors were harvested for downstream analysis.

For xenograft experiments, 5 × 10^6^ OCI-Ly18 cells were mixed at a ratio of 1:1 in Matrigel (Corning, #356237) and subcutaneously injected into the right flank of NSG mice. Two intravenous doses of CAR T-cells were administered to tumor-bearing mice on day 12 and day 17 post-tumor implantation. When tumors reached approximately 2000 mm^3^, mice were euthanized, and tumors were harvested for downstream analysis.

### Flow cytometry analysis

Tissue immune profiling was carried out similarly to that described previously ^43, 44^. Briefly, tumor tissue was dissected and digested with 1 mg/mL Collagenase D (Roche, #11088882001) and 100 mg/mL DNase I (Roche, #10104159001) in RPMI medium supplemented with 2% FBS for 40 minutes with continuous agitation at 37°C, 200 RPM. The digestion mixture was passed through a 100 μm cell strainer (Corning, #431752) to prepare a single-cell suspension and then was washed with FACS solution (PBS supplemented with 2 mM EDTA and 1% FBS). After that, single cells were incubated with the surface, followed by intracellular staining antibodies if required. All conjugated antibodies were diluted to 1:100-1:200 in FACs buffer before use and incubated with cells at 4°C for 30 mins. Following staining, cells were washed with PBS and subjected to flow cytometry analysis.

Splenocytes were collected from the spleen of experimental animals. The spleen was dissected, mashed through a 70 μm cell strainer (Corning, #87712), and treated with red blood cell (RBC) lysis buffer to remove erythrocytes. After that, the single cells were processed in a manner similar to that of tumor samples.

For CD8^+^ T-cell cytokine production analysis, 2 × 10^6^ cells were treated with 2μL of activation cocktail/mL in RPMI-1640 Medium (Gibco, #22400089), supplement with 10% HyClone fetal bovine serum (Cytiva, #SH30071.03), 1mM sodium pyruvate (Gibco, #11360070),1× Antibiotic/Antimycotic Solution(Cytiva, #SV3007901), 55μM 2-mercaptoethanol (Sigma, #60242), for 4-6 hours before collection for the staining process.

### Immunofluorescence staining

For immunofluorescence, the OCT-stored tissues were cut into 8.0 μm sections onto glass slides and fixed in a 4% PFA solution (Thermo, #J19943-K2) at room temperature for 15 minutes. The tissues were washed twice with PBS. The samples were incubated in 0.1% Triton in PBS for 10 min at RT, followed by blocking in PBS supplemented with 0.1% Triton and 5% donkey serum (Sigma, #S30-100ML). Tissue samples were incubated with primary antibodies overnight at 4 °C and washed three times with PBS-T before being incubated with respective secondary antibodies at room temperature (RT) for 1 hour. ProLong Gold Antifade Reagent with DAPI (Invitrogen, #P36931) was added. The images were taken using the Leica DM6000 microscope. The quantification of immunofluorescence intensity was performed by ImageJ. Three fields in each section were quantified.

The following primary antibodies were purchased and used in the experiments: Rabbit anti-CD8a (Thermo Fisher, #MA5-29682; dilution 1:200), Rabbit anti-fibronectin (FN) (Sigma, #F3648; dilution 1:400), Mouse anti-Collagen I (Col-I) (Thermo Fisher, #MA1-26771; dilution 1:2000), Rabbit anti-Fibroblast Activation Protein (FAP) (Abcam, #Ab53066; dilution 1:300), Rat anti-CD31 (BD Biosciences, #553370; dilution 1:1000)

The following secondary antibodies were used: Donkey anti-rabbit 488 (Biolegend, #406416, dilution 1:800), Donkey anti-Mouse 555 (Thermo, #A21209, dilution 1:500), Donkey anti-Rat 594 (Thermo, #A31570, dilution 1:500)

### Tumor-infiltrating lymphocytes (TILs) and splenic T-cell isolation

For TILs isolation, MC38 (5×10^5^ cells/mouse) were injected s.c. into WT mice. 21 days after injection, tumors were collected. The tissues were then cut into small pieces using dissection scissors, followed by incubation in a digestion solution containing 1 mg/ml Collagenase D (Roche, #11088882001) and 100 µg/ml DNase I (Roche, #10104159001) for 40 minutes with continuous agitation (200 RPM) at 37°C. Digestion was halted by adding pre-warmed FBS and passing it through a 100 μm strainer (Corning, #431752) to obtain a single-cell suspension, followed by centrifugation at 500×g for 5 minutes. The cell pellet was resuspended in 5 mL of RBC lysis buffer at room temperature for 5 minutes. The cells were then washed twice with PBS. TILs were isolated using Percoll (Cytiva, #45-001-747) density gradient centrifugation. Briefly, 9 mL of 44% Percoll solution was added to a 50 mL centrifuge tube, followed by slow underlaying of 6 mL of 66% Percoll. The single-cell suspension from the tumor tissue was then placed on top of the upper layer, and centrifugation was performed at 800 ×g at 4°C for 30 minutes. After centrifugation, the interphase between the two Percoll layers, which contained the TILs, was collected and washed twice with PBS. The TILs were then processed for T-cell isolation using the EasySep™ Mouse T-cell Isolation Kit (STEMCELL, #19851), following the manufacturer’s instructions.

Spleen-derived T-cells were isolated by harvesting the spleen and mechanically disrupting it through a 70 μm strainer (Corning, #87712) to obtain a single-cell suspension. The suspension was treated with RBC lysis buffer to eliminate erythrocytes. The cells were then washed and processed for T-cell isolation using the EasySep™ Mouse T-cell Isolation Kit (STEMCELL, #19851), according to the manufacturer’s instructions.

### Quantitative Polymerase Chain Reaction (qPCR)

Cell pellets were collected for RNA isolation using the RNeasy Plus Mini Kit (QIAGEN, #74134). The mRNA was then converted to cDNA via reverse transcription (RT) using the High-Capacity RNA-to-cDNA Kit (Applied Biosystems, #4388950). Subsequently, qPCR was performed using specific primers for SPTBN2 and GAPDH, along with PowerUp SYBR Master Mix (Applied Biosystems, #A25777) on a QuantStudio 3 system. The ΔCT value was calculated by subtracting the CT value for SPTBN2 from the CT value for the endogenous control (GAPDH). The expression levels were represented as the relative change between groups.

### Western blot analysis

Human CAR T-cells were collected, counted, and stored at −80°C until further use. All samples were adjusted to the same concentration in the sample buffer. Immunoblotting was performed as previously described ^40^. The following primary antibodies were used: anti-β-III Spectrin (Life Technologies, #711584; dilution 1:500) and anti-Vinculin (Cell Signaling Technology, #4650S; dilution 1:10,000). HRP-conjugated goat anti-rabbit IgG (Millipore, #AP187P; dilution 1:10,000) or HRP-conjugated goat anti-mouse IgG (Cell Signaling Technology, #7076S; dilution 1:10,000) was used as the appropriate secondary antibody. Band intensities were quantified using ImageJ, and the β-III Spectrin to Vinculin ratio was calculated and normalized to the WT control.

### Gene expression correlation

Using TIMER 2.0 (https://compbio.cn/timer2/, accessed May 2025), we analyzed the correlation of CD8A gene expression across various cancer types from the TCGA dataset. The “Correlation” module was used to generate expression scatterplots between CD8A and SPTBN2 gene within specific cancer types, providing Spearman correlation coefficients and corresponding statistical significance. In addition, CD8^+^ T-cell infiltration was assessed using the QUANTISEQ algorithm to evaluate the relationship between gene expression and the levels of immune cell infiltration.

### Survival correlation

We evaluated the association between SPTBN2 mRNA expression and patient survival using the KMplot database (https://kmplot.com; accessed March 2026). This platform enables survival analysis across more than 40,000 samples spanning 21 tumor types, based on gene expression levels. In the resulting Kaplan– Meier plots, the x-axis represents survival time (in months), while the y-axis indicates the probability of survival.

### Correlation with B cell aplasia

Processed single-cell data reported in ^38^ were downloaded from NCBI Gene Expression Omnibus under accession number GSE262072. Using R v4.5.1 and Seruat v5.2.0, the provided counts data was split by sample and re-normalized using default parameters using the NormalizeData function in Seurat. Subsequent statistical analyses between relevant clinical groups were performed using Wilcox.test.

### Quantification and statistical analysis

All animal experiments described here are representative of at least five mice for each group unless specifically indicated. For in vitro experiments, the samples were processed at least in triplicate. All data here were shown as average ±S.E.M. Statistical analysis between two groups was conducted with a 2-tailed Student’s T-test, and multiple comparisons were performed by using a one-way ANOVA followed by a Tukey post-hoc test with P values <0.05 considered significant. All statistical analyses were performed using GraphPad Prism 10, and the data for each experiment are clearly described in their respective figures and figure legends.

## SUPPLEMENTARY DATA

## LEGENDS TO SUPPLEMENTARY FIGURES

**Figure S1.**
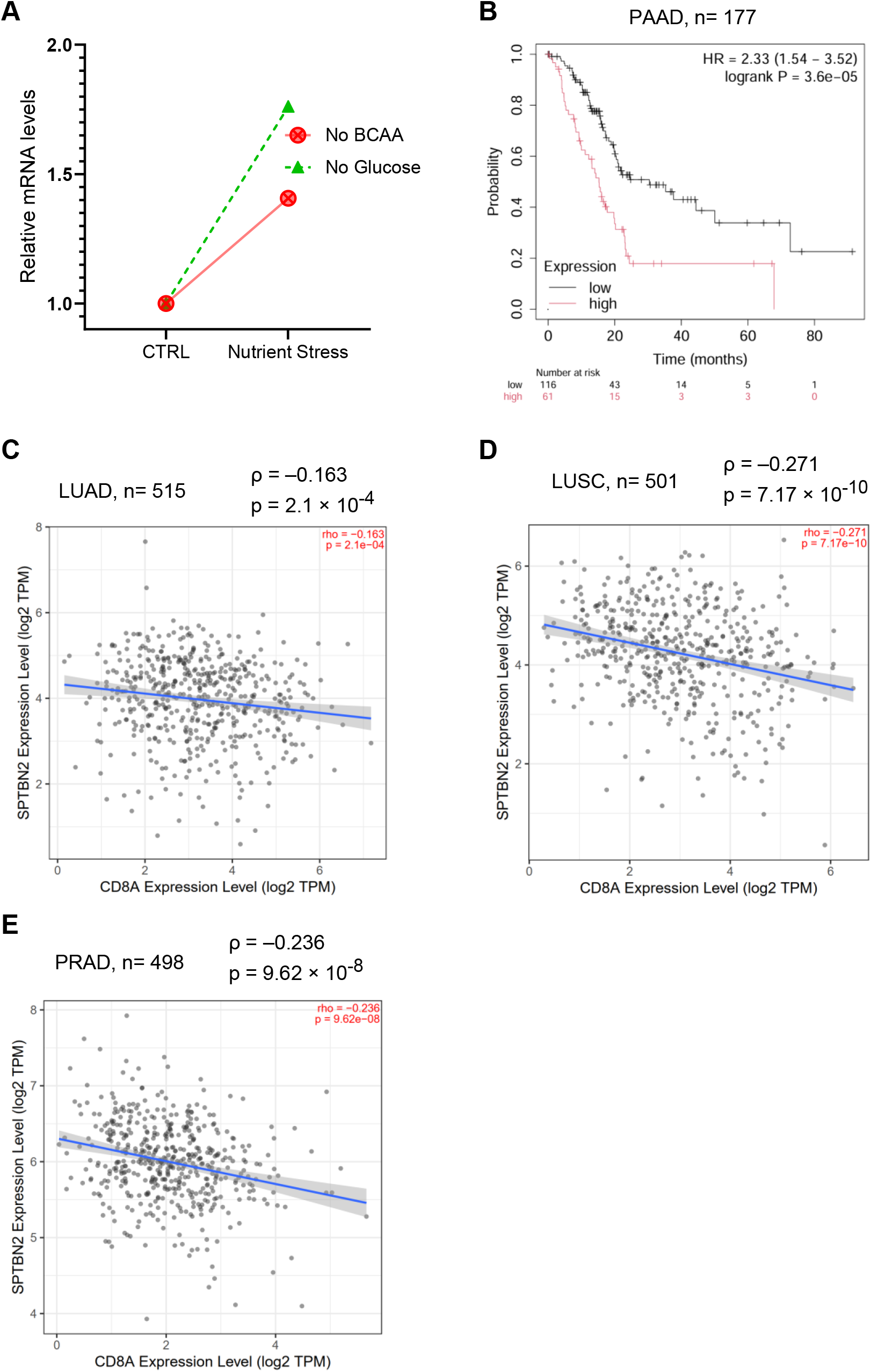
SPTBN2 is up regulated in the tumor microenvironment. A. Normalized levels of total *SPTBN2* mRNA in CD8^+^ T-cells cultured without glucose or branched chain amino acids (BCAA). Data is presented as a fold change relative to the control group (complete media); n=3. B. Association between *SPTBN2* expression and overall survival in human PAAD (n=177) samples, analyzed using the KMplot database. Patients were stratified into high and low expression groups based on the median SPTBN2 expression. The hazard ratio (HR), log-rank p value, and 95% confidence interval (Cl) are shown. C. Correlation of expression levels for CD8A and SPTBN2 in human lung adenocarcinoma (n=515) samples from TCGA. D. Correlation of expression levels for CD8A and SPTBN2 in human lung squamous cell carcinoma (n=501) samples from TCGA. E. Correlation of expression levels for CD8Aand SPTBN2 inhuman prostate adenocarcinoma (n=498) samples from TCGA. All analyses were performed using the TIMER2.0 platform, and Spearman correlation coefficients and p-values are indicated.

**Figure S2.**
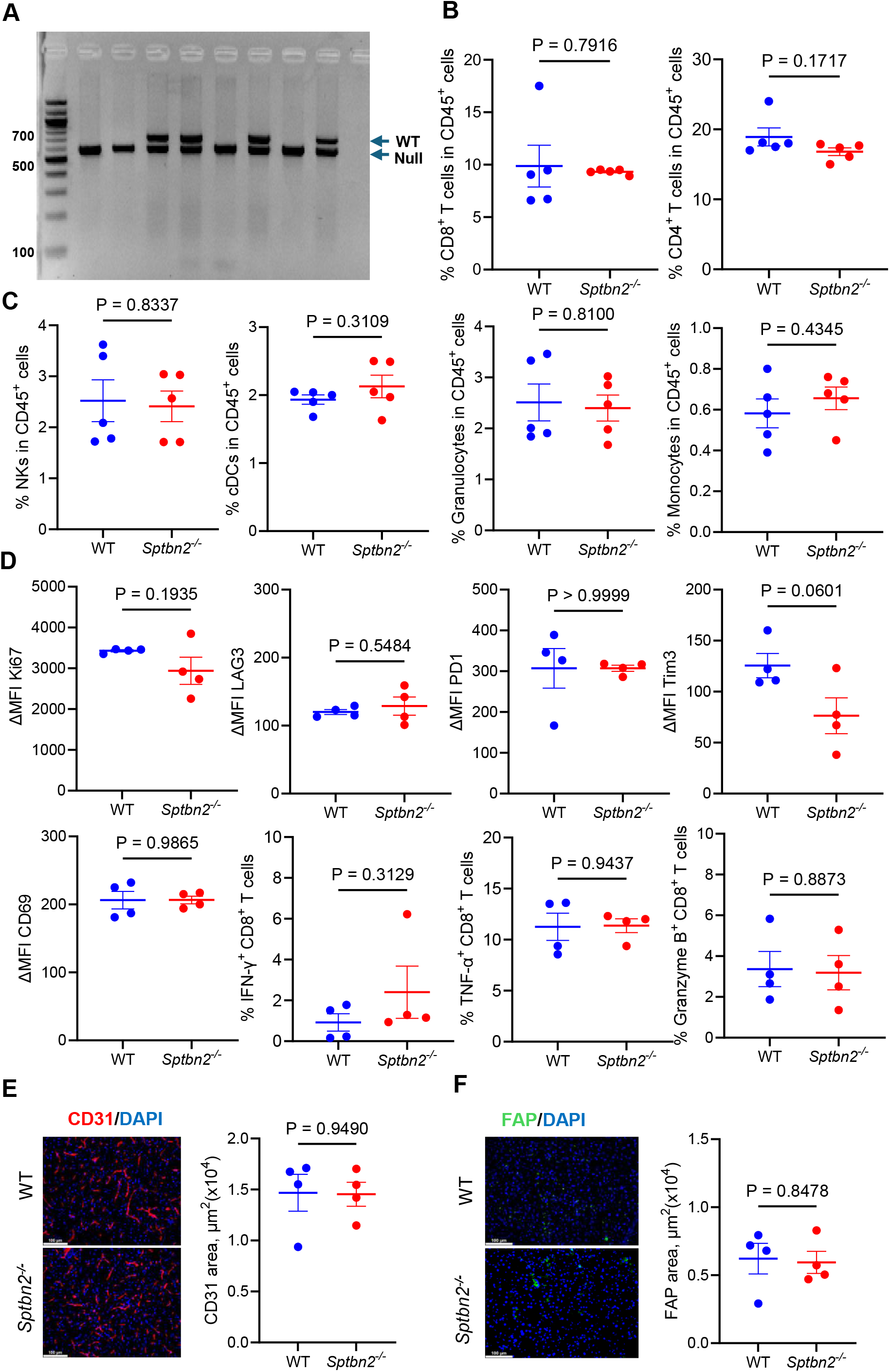
SPTBN2 supports intratumoral angiogenesis and stromagenesis. A. PCR-based genotyping analysis of DNA from the tails of WT and Sptbn2^-/-^ mice. B. Frequencies (% in live CD45^+^) of CD8^+^ and CD4^+^ T-cells in the spleen of WT and *Sptbn2*^-/-^mice (n=5). C. Frequencies (% in live CD45^+^) of NKs, eDCs, Monocytes, and Granulocytes in the spleen of WT and *Sptbn2*^*-/-*^ mice (n=5). D. Flow cytometry analysis of MTI of Ki67, LAG3^+^, PD1^+^, Tim3^+^, CD69^+^ and frequencies of IFN-α^+^, TNF-α^+^, Granzyme B^+^ of CD8^+^T-cells in the spleen of WT and *Sptbn2*^-/-^ mice (n=4). E. Representative immunofluorescence images showing CD31 staining in normal WT and *Sptbn2*^*-*/-^ pancreata (n=4). The quantification shows the percentage of CD31 positive areas. Scale bar: 100 pm. F. Representative immunofluorescence images showing TAP staining in normal WT and *Sptbn2*^-/-^ pancreata (n=4). The quantification shows the percentage of TAP positive areas. Scale bar: 100 pm. Data are presented as meant SEM. Statistical comparisons were conducted using the Student’s t-test.

**Figure S3.**
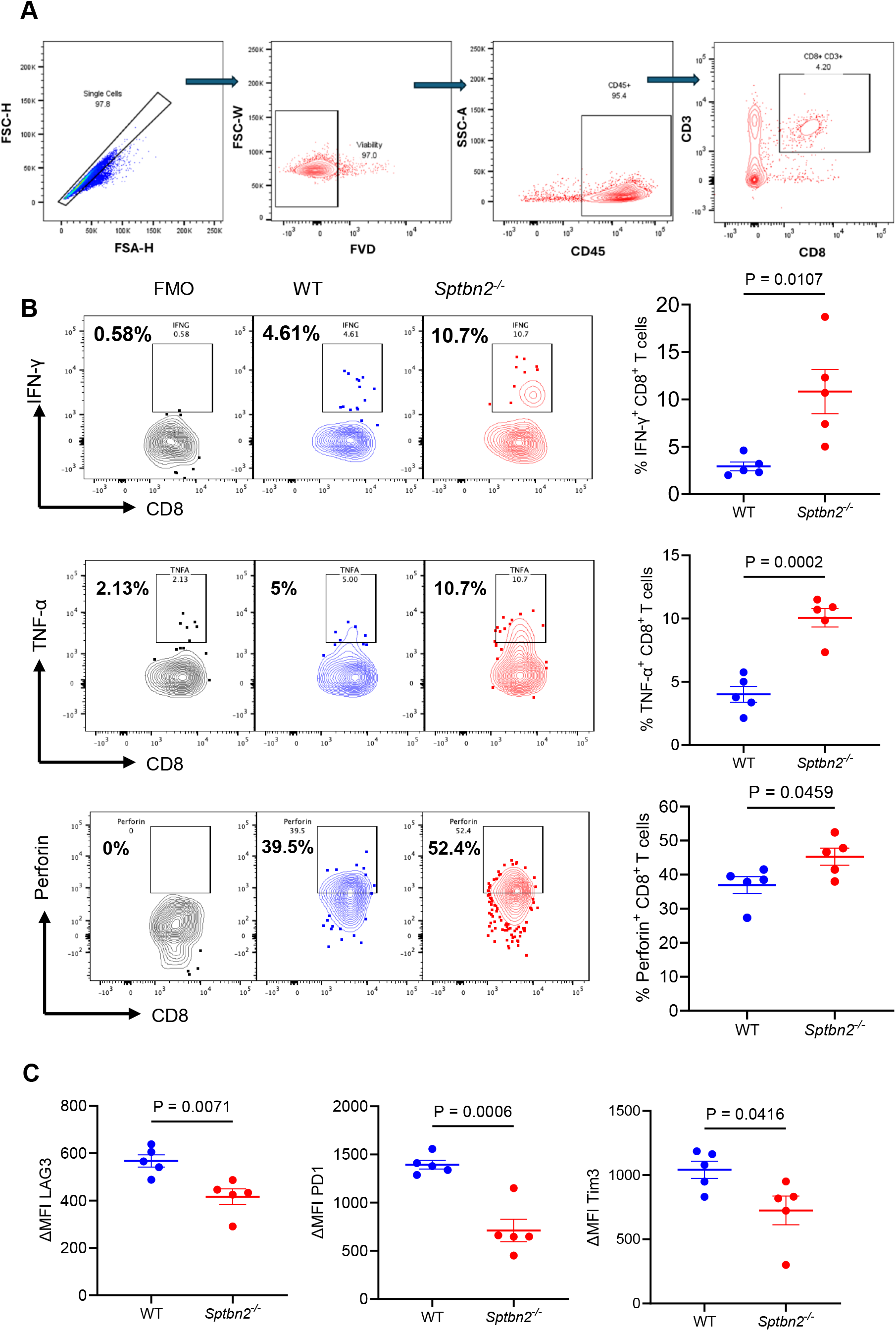
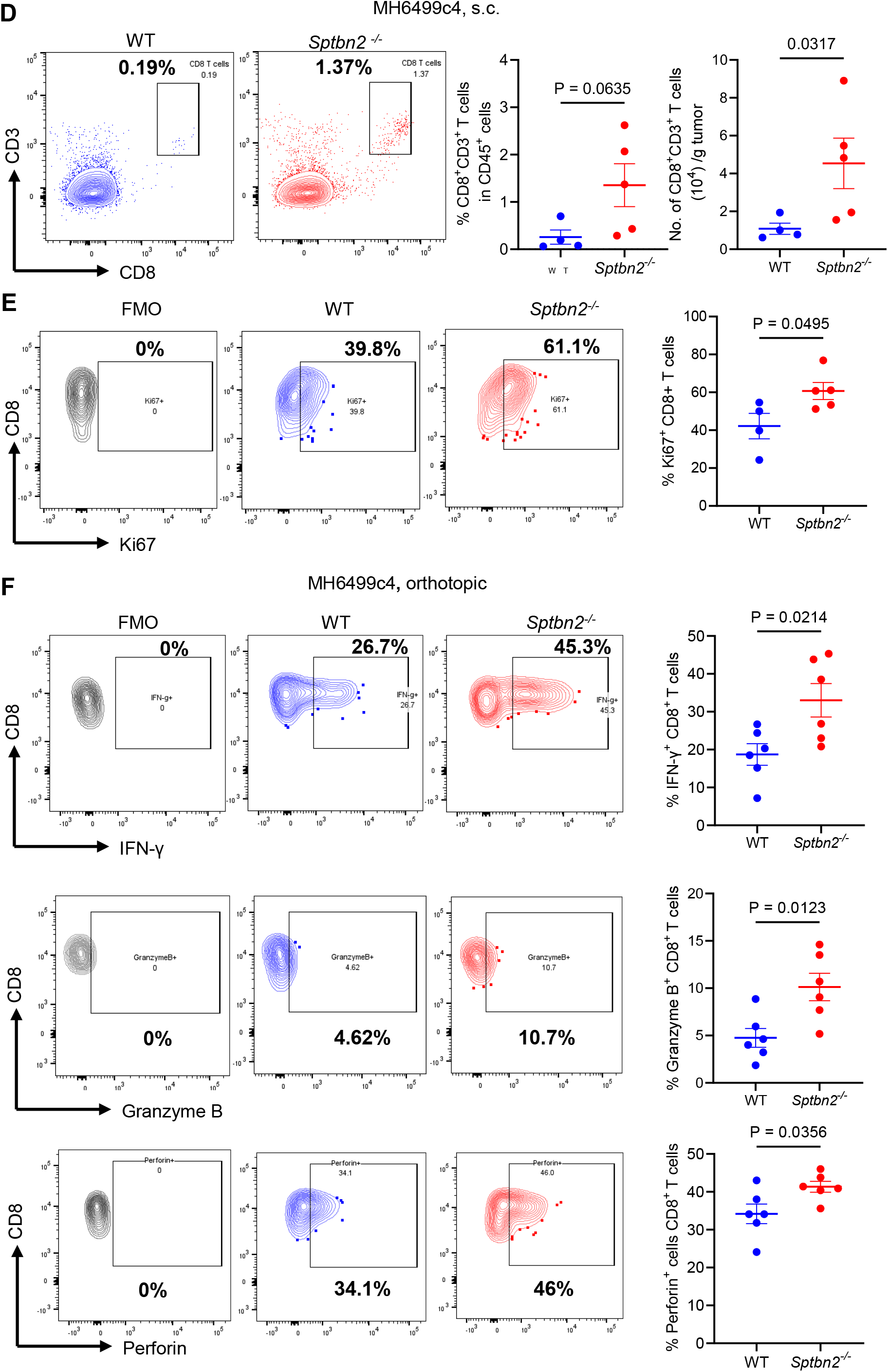
SPTBN2 inactivates tumor-infiltrating CTLs. A. Gating strategy for immune profiling of tumor/splenic tissues. B. Frequencies of IFN-γ^+^, TNF-*α*^+^, Perforin^+^ CD8^+^ T-cells isolated from the subcutaneous MH6419c5 tumors grown in WT or Sptbn2^/^ mice (n=5). C. Frequencies of LAG3^+^, Tim3^+^, and PD-1^+^ CD8^+^ T-cells from the subcutaneous MH6419c5 tumors grown in WT or *Sptbn2*^-/-^ mice (n=5). D. Frequency and number of CD8^+^ T-cells from the subcutaneous MH6499c4 tumors grown in WT or *Sptbn2*^*-/-*^ mice (n=4-5). E. Frequency of Ki67^+^ CD8^+^T-cells from the subcutaneous MH6499c4tumors grown in WT or *Sptbn2* ^-/-^ mice (n=4-5). F. Frequencies of IFN-γ^+^, Granzyme B’, Perforin’ CD8’T-cells from the orthotopic MH6499c4 tumors grown in WT or Sptbn2^-/-^ mice (n=6). Data are presented as mean ± SEM. Statistical comparisons were conducted using the Student’s t-test.

**Figure S4.**
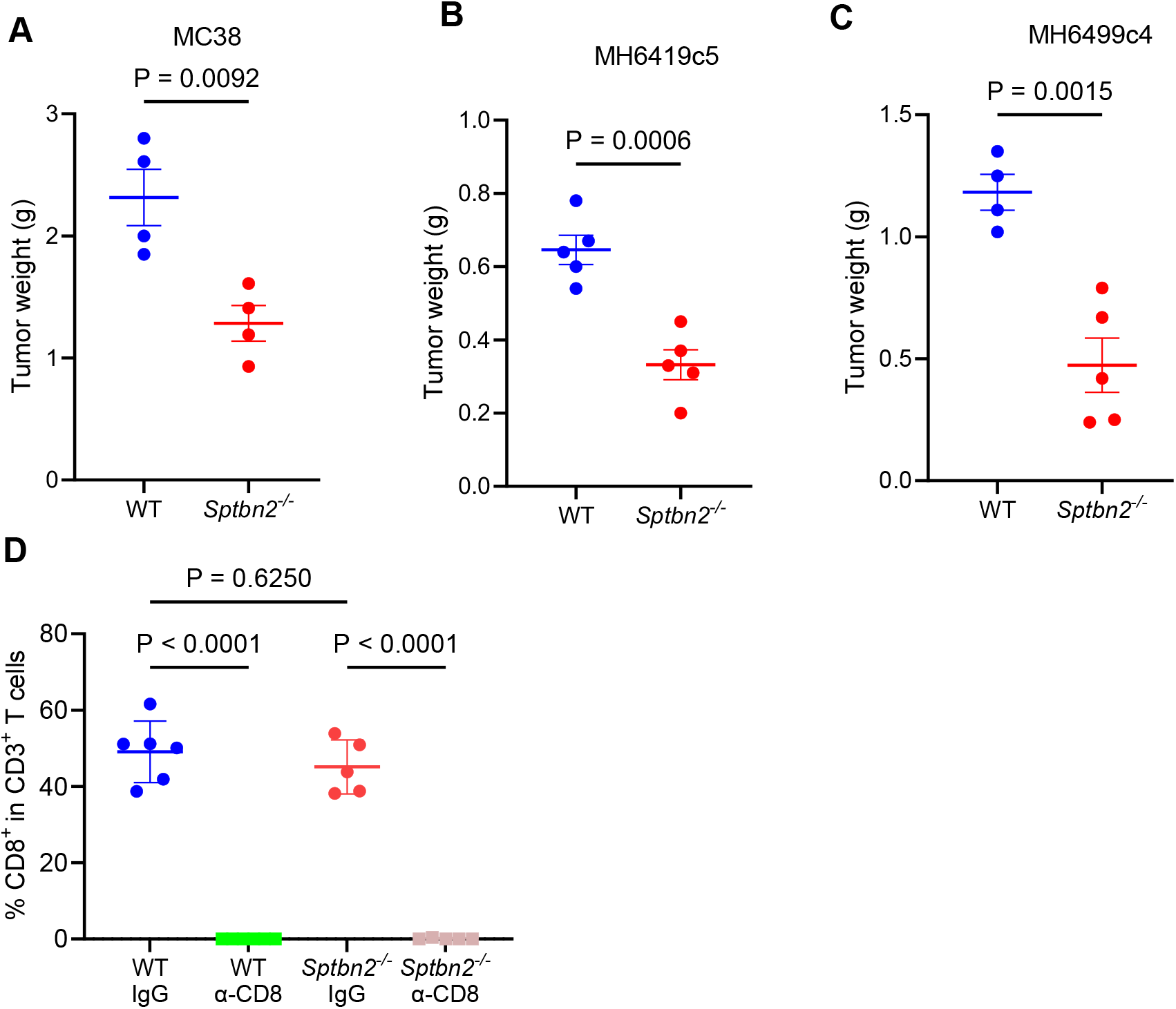
SPTBN2 promotes tumor growth in a CTL-dependent manner. A. Tumor masses of subcutaneous MC38 in WT and *Sptbn2*^-/-^ mice (n=4). B. Tumor masses of subcutaneous MH6419c5 in WT and *Sptbn2*^-/-^ mice (n=5). C. Tumor masses of subcutaneous MH6499c4 in WT and *Sptbn2*^-/-^ mice (n=4-5). D. Percentage of CD3^+^CD8^+^ T-cells in the peripheral blood of tumor-bearing mice treated with anti-CD8 antibodies. Data are presented as mean ± SEM. Statistical comparisons were conducted using the Student’s t-test for panels AC and one-way AN OVA followed by Tukey’s post hoc test for panel D.

**Figure S5.**
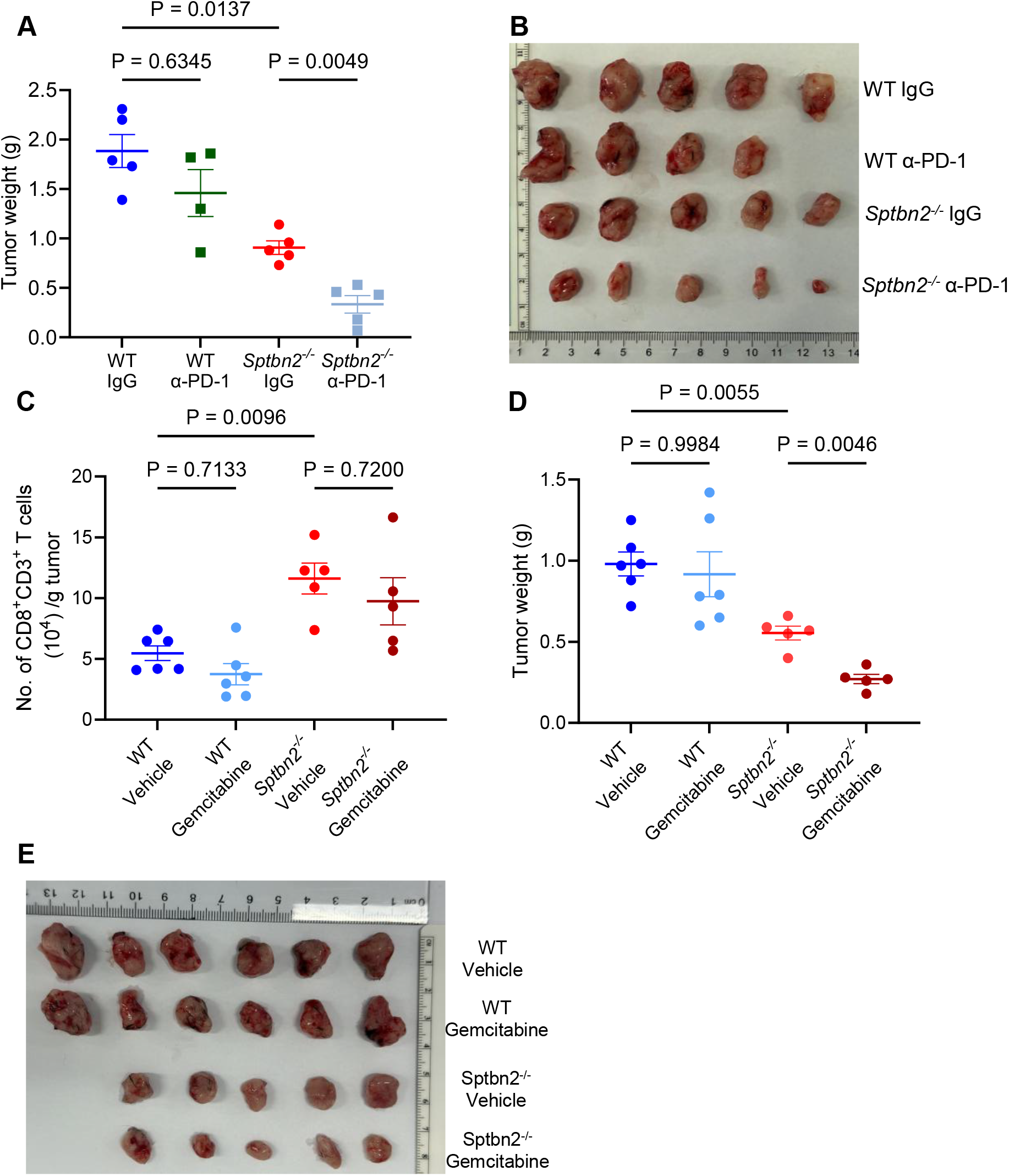
SPTBN2 confers PDAC resistance to chemotherapy and immune checkpoint blockade. A. Tumor weight of subcutaneous MH6419c5 tumors in WT and *Sptbn2*^-/-^ mice treated with α-PD1 or IgG, as described in **5A**. B. Tumor image of subcutaneous MH6419c5 tumors in WT and *Sptbn2*^-/-^ mice treated with α-PD1 or IgG, as described in **5A**. C. Number of CD8^+^ T cells per gram of subcutaneous MH6419c5 tumors in WT and *Sptbn2*^-/-^ mice treated with Gemcitabine or vehicle, as described in **5F**. D. Tumor weight of subcutaneous MH6419c5 tumors in WT and *Sptbn2*^-/-^ mice treated with Gemcitabine or vehicle, as described in **5F**. E. Tumor image of subcutaneous MH6419c5 tumors growth in WT and *Sptbn2*^-/-^ mice treated with Gemcitabine or vehicle, as described in 5**F**. Data are presented as mean ± SEM. Statistical comparisons were conducted using one-way ANOVA followed by a Tukey post hoc test.

**Figure S6.**
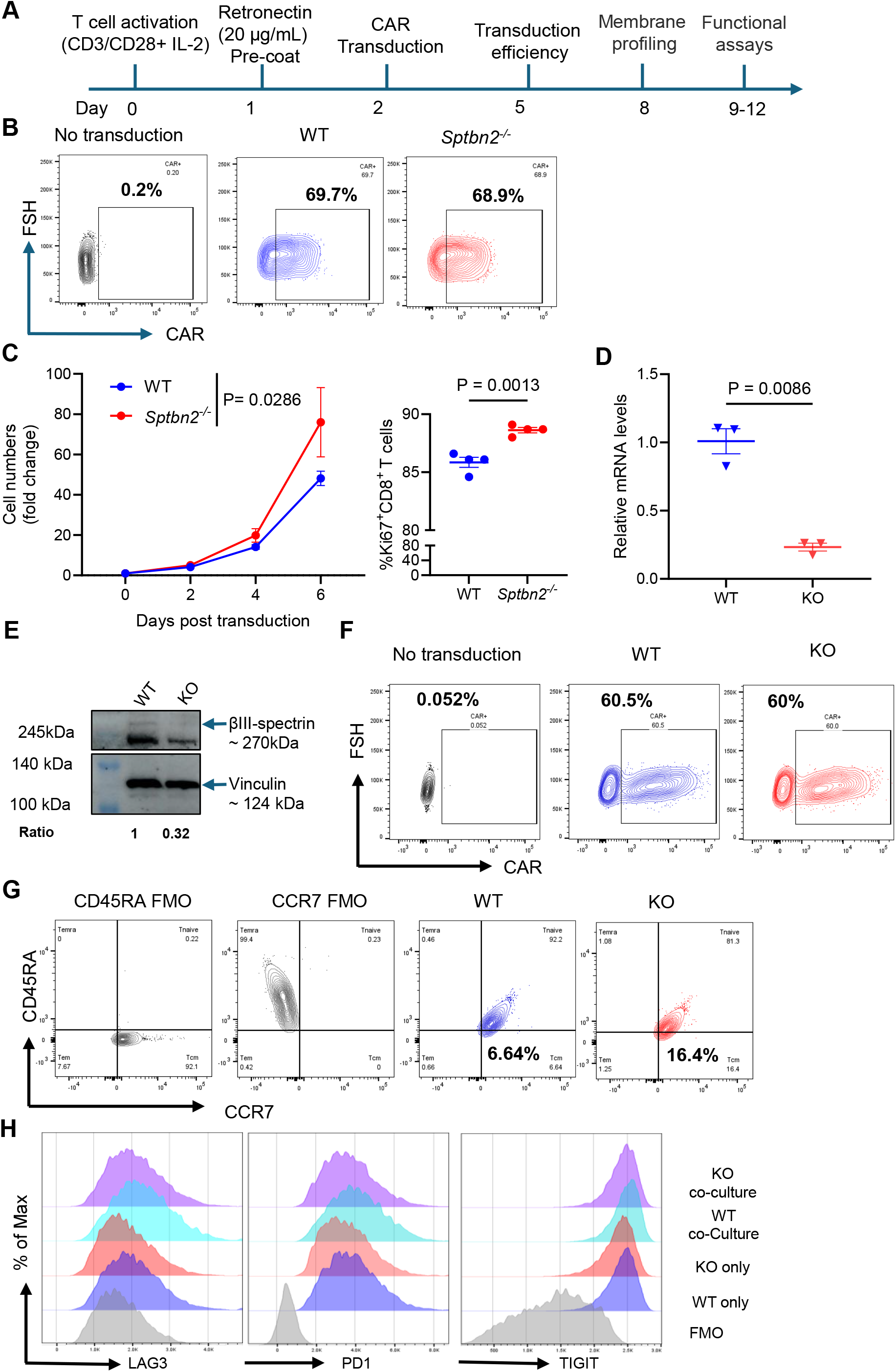
Expression of SPTBN2 in CAR T-cells promotes their inactivation and in vitro exhaustion. A. Schematic representation of anti-hCD19 CAR T-cells derived from WT and *Sptbn2*^-/-^ splenocytes, showing their generation, quality control, and subsequent experimental applications. B. Representative flow cytometry analysis of CAR expression in mouse T-cells of the indicated genotypes after hCD19-BBztransduction C. The *in vitro* expansion of WT or *Sptbn2*^-/-^ CAR T-cells, from independent CAR T-cell batches (n =4). The data represents the fold change in cell number relativeto the initial seeding point. Right panel shows frequency of the Ki67+ cells in these populations. D. qPCR analysis of *SPTBN2* mRNA levels in WT or SPTBN2 KO human CAR T-cells E. Western blot analysis of the SPTBN2 (βlll spectrin) levels in WT or CRIPSR generated SPTBN2 KO human CAR T-cells. Vinculin was used as a loading control. F. Representative flow cytometry analysis of CAR expression in human T-cells of the indicated genotypes G. Representative flow cytometry analysis of central memory (CD45RA ^−^CCR7^+^) in WT or SPTBN2 KO human CD8^+^ CART-cells (results quantified in **Figure 6E**). H. Representative flow cytometry analyses of MFI of LAG3, PD-1, and TIGIT on CD8^+^ CAR T-cells cultured with or without OCI-Lyl8 tumor cells at an effector-to-target (E: T) ratio of 5:1 for 20 hr (results quantified in **Figure 6F**). Data are presented as mean± SEM. Statistical comparisons were conducted usingthe Student’s t-test or one-way ANOVA followed by a Tukey post hoc test.

**Figure S7.**
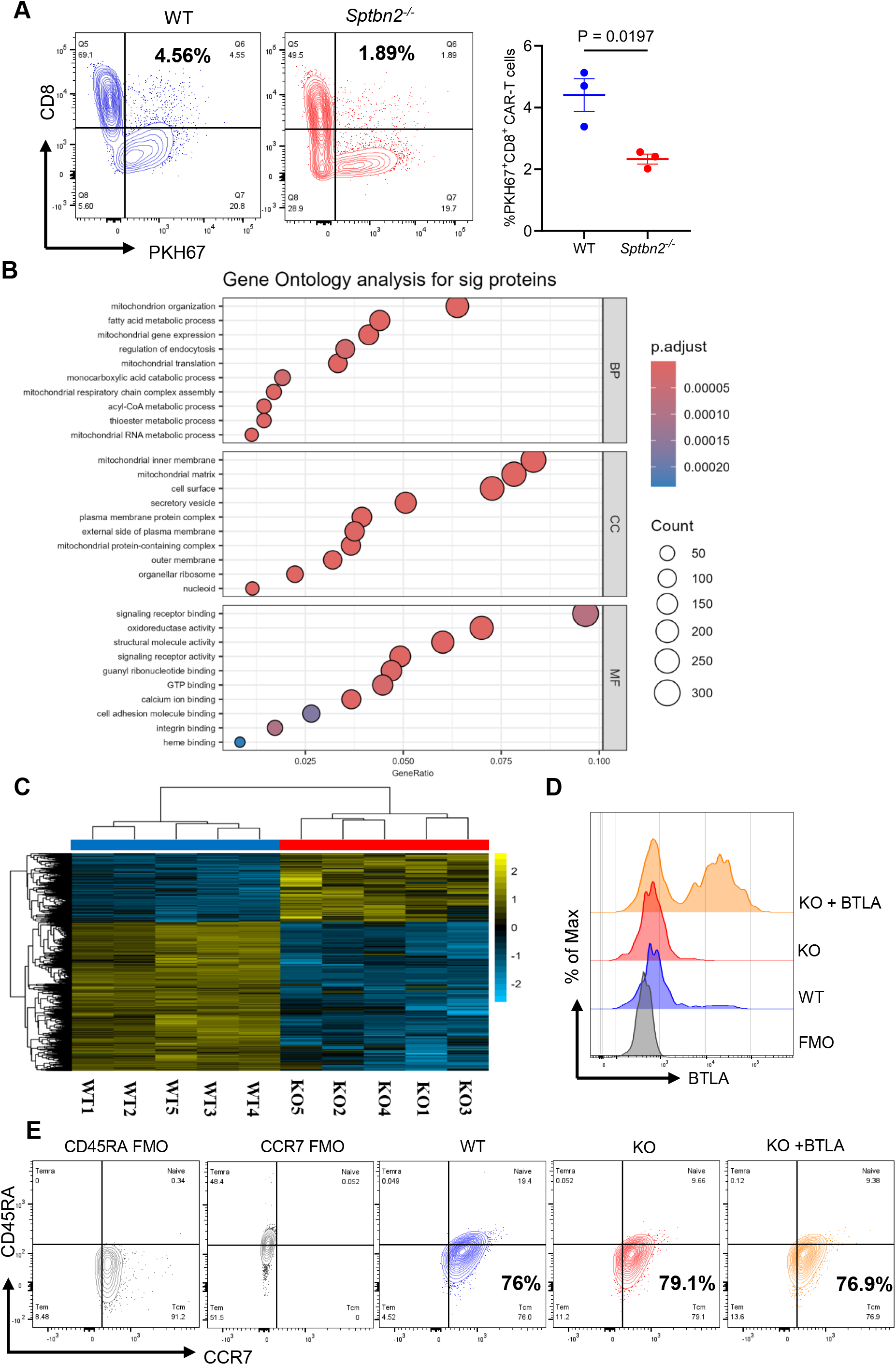
SPTBN2 regulates cell surface levels of plasma membrane proteins in CAR T-cells. A. Analysis of the lipid membrane dye (PKH67) transfer from B16F10-hCD19 tumor cells to CAR T-cells derived from WT or *Sptbn2*^-/-^ mice (n=3). CAR T-cells were co-cultured with B16F10.hCD19 cells at an E: T ratio of 5:1 for 15 minutes. The quantification represents the percentage of PKH67^+^ CD8^+^ CAR T-cells. B. Gene Ontology (GO) enrichment plot of significantly altered proteins between WT and *Sptbn2*^-/-^ CAR T-cells (n = 5). C. Heat map analysis of significantly altered proteins between WT and *Sptbn2*^-/-^ CAR T-cells (n = 5). D. Representative flow cytometry analysis of BTLA levels on indicated human CART-cells E. Representative flow cytometry analysis of central memory (CD45RA ^−^CCR7^+^) markers on indicated human CART-cells (quantified in Figure **7E**). Data are presented as mean ±SEM. Statistical comparisons were conducted using the Student’s t-test.

**Figure S8.**
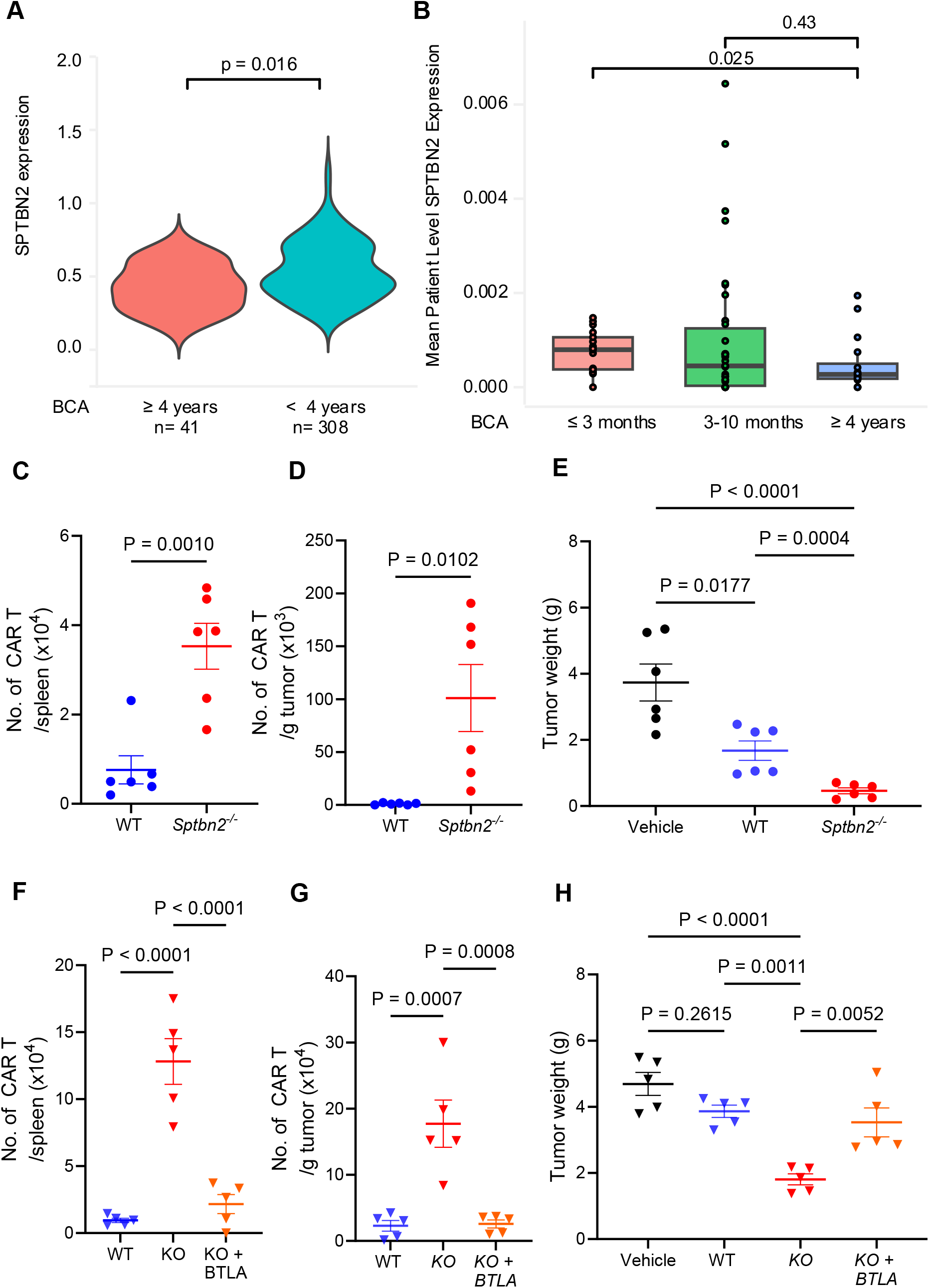
Inactivation of SPTBN2 in CAR T-cells supports their persistence and enhances therapeutic efficacy. A. Violin plot of normalized SPTBN2 expression in CD19-stimulated manufactured CAR T cells stratified by simplified patient BCA groups. P-values shown are unadjusted. B. Boxplot of normalized mean SPTBN2 expression by patient in manufactured CART cells stratified by BCA outcome ^1^. Each dot represents a patient. Average patient level expression of SPTBN2 in manufactured CAR T cells cultured in the basal condition are shown. P-values shown are unadjusted. C. Number of mouse CART-cells in the spleens of tumor-bearing mice collected at the experimental endpoint described in Figure **8B**. D. Quantification of mouse CAR T-cells per gram of tumor tissue collected at the experimental endpoint, as described in Figure **8B**. E. Tumor weight at the endpoint of B16F10.hCD19 tumor growth in Rag1^-/-^ mice treated with WT or *Sptbn2*^*-/-*^ CAR T, as described in Figure **8B**. F. Number of human CAR T-cells in the spleen of NSG tumor-bearing mice collected at the experimental endpoint described in Figure **8E**. G. Quantification of human CAR T-cells per gram of tumor tissue collected at the experimental endpoint, as described in Figure **8E**. H. Tumor weight at the endpoint of OCI-Lyl8 tumors growth in NSG mice treated with WT or SPTBN2 KO CAR T-cells, as described in Figure **8E**. Data are presented as mean± SEM. Statistical comparisons were conducted using a two-sided unpaired Wilcoxon rank sum test for panels A-B, Student’s t-test for panels C-D, and one-way ANOVA followed by Tuke’s post hoc test for panels E-H. 1. Bai, Z. *et al*. Single-cell CAR T atlas reveals type 2 function in 8-year leukaemia remission. Nature **634**, 702-711 (2024).

